# Does the paradigm of genotype-environment associations need to be re-assessed? The paradox of adaptive phenotypic clines with non-clinal patterns in causal alleles

**DOI:** 10.1101/2022.08.03.502621

**Authors:** Katie E Lotterhos

## Abstract

Multivariate climate change presents an urgent need to understand how species adapt to complex environments. Population genetic theory predicts that loci under selection will form monotonic allele frequency clines with their selective environment, which has led to the wide use of genotype-environment associations (GEAs). This study elucidates the conditions under which allele frequency clines are more or less likely to evolve as multiple quantitative traits adapt to a multivariate environment. A novel set of simulations was created that all evolved similar phenotypic clines, but with varying proportions of causal alleles with clines. Phenotypic clines evolved mostly without clines in the causal allele frequencies under conditions that promoted unique combinations of mutations to achieve the multivariate optimum in different parts of the landscape. Although univariate and multivariate GEA methods failed to accurately infer the genetic basis of adaptation under a range of scenarios, individual multivariate traits could be accurately predicted from genotype and environmental data without any knowledge of the genetic architecture. This research challenges the utility of GEAs for understanding the genetic basis of adaptation to the environment, and instead suggests that multivariate trait predictions are a more fruitful approach for genomic forecasting and assisted gene flow efforts.

## Introduction

Clines have a rich history of study in biology. Clines were first described as a gradient in a measurable character (1), but they can also be a gradient in a genotype or allele frequency. Prior to the 1950’s, geographic differentiation and speciation were thought to require some kind of isolation (2). Starting around the 1950s, however, population geneticists became more interested in the interaction between gene flow and selection in continuous populations along environmental gradients (2–4). Population genetic theory that predicts clines in allele frequency along an environmental gradient arises from the assumption that the fitness of one allele increases along an environmental gradient, while the other decreases (2,5–8).

As a result of this theory, ecological geneticists have long searched for clinal patterns in allele frequency along environmental gradients, dating back to Dobzhansky’s work on clines in *Drosophila* chromosome inversions (9,10), followed by many examples in allozymes (e.g. (11–15)). With recent advances in sequencing, the search for allele frequency clines has extended to the entire genome, with a focus on understanding the genetic basis of adaptation to the environment (16–18). Many statistical approaches have been developed to determine if associations between allele frequency and an environmental gradient are significant after correcting for neutral population structure; these methods are referred to “genotype-environment associations” (hereafter GEA, sensu 19), “genetic environment associations” (also GEA, sensu 17, 20), or “environment-allele associations” (16). Verbal arguments have claimed that beneficial variants with weak phenotypic effects will lead to subtle shifts in allele frequency that correlate with environmental variables (21–23). In addition, multilocus models of polygenic adaptation across an environmental gradient predict that phenotypic clines will evolve by a series of staggered allele frequency clines (24–26).

GEA methods are based on the distinct concept that adaptive alleles will form clines with their respective selective environments. Since 2010, the most widely used methods (27–32), reviews (16,17,26,33), methods evaluations (34–39), and high-profile applications of GEA methods in different taxa (23,40–44) have been cited over 7700 times (Google Scholar, July 2022). A scientific paradigm is a distinct set of concepts or thought patterns, including theories, research methods, postulates, and standards for what constitutes legitimate contributions to a field. The GEA concept and methods meet this definition of a paradigm, and more specifically Kuhn’s definition of a “local” scientific paradigm (i.e., a typical example or exemplar) for how to study the genetic basis of adaptation to the environment (45,46). Although other lines of inquiry such as *F_ST_* outlier tests for genetic differentiation are also used, *F_ST_* outlier tests do not explicitly link genetic differences to the environment (17).

GEA methods have almost exclusively been tested against data simulated under the assumption that the fitness of one allele increases along an environmental gradient while the other decreases, which evolves allele frequency associations with the climate variable (16,27,29–31,34–37). This practice may have led to overly optimistic performance of GEA methods if the evolution of local adaptation does not lead to the evolution of clines in the causal allele frequencies (the first principles issue). Secondly, even when clines in causal allele frequencies do evolve, GEAs may fail to appropriately correct for structure or over-correct for structure, leading to a failure to accurately detect the genetic basis of adaptation (the statistical issue). An extension of the statistical issue is the question of whether multivariate ordination can be used to accurately infer multivariate adaptation. If these issues prove to be relevant, then reliance on the study of clines may hinder our ability to accurately detect the genetic basis of adaptation and translate landscape/seascape genomic data into a reliable prediction of populations’ vulnerability to climate change.

The goals of this study were to broadly understand (i) the conditions under which allele frequency clines are more or less likely to evolve as quantitative traits adapt to complex environments, (ii) the ability of univariate (correlation, latent factor mixed models) and multivariate (redundancy analysis) GEA methods that do or do not correct for structure to correctly infer the genetic basis of adaptation, and (iii) whether useful predictions regarding local adaptation could be made from genome and environmental data without knowledge of the genetic architecture. To achieve these goals, I created a novel set of quantitative genetic simulations that were designed to advance our understanding of how multivariate adaptation in complex environments can result in non-intuitive evolutionary dynamics. The simulations included (i) up to two quantitative traits, each of which adapted to a different environmental pattern, (ii) genetic architectures ranging from oligogenic (a few loci with large effects on the trait) to highly polygenic (thousands of loci with individually small effects), (iii) pleiotropic or non-pleiotropic effects of mutations on multiple traits, and (iv) complex landscapes and demographies. Alleles had additive effects on the quantitative traits, which were under stabilizing selection within populations but the optimum trait value was linearly associated with the selective environment. Across all simulations, clines evolved between the quantitative trait and the deme environment, but the proportion of quantitative trait nucleotides (QTNs) that evolved clines depended on the parameters.

## Results

I conducted replicate forward-time simulations of a metapopulation adapting to a heterogeneous spatial environment with SLiM v. 3.6 (47,48) to create single nucleotide polymorphism data for each individual. I simulated 225 parameter levels (15 demographies x 15 genetic architectures) of 10 replicates each, for a total of 2250 simulations. The levels varied in (i) the relationship between the landscape and the environmental trait optimum (with cases in which geographic distance did or did not correspond to environmental or genetic distance: *Stepping Stone Clines, Stepping Stone Mountain, Estuary Clines* **Figure 1A),** (ii) the demography described by migration rates and effective population size (with cases that confounded genetic drift with selection and cases that confounded population structure with selection, **Figure 1B),** (iii) the genic level of both traits (with cases spanning from oligogenic to highly polygenic, **Figure 1C),** and (iv) the pleiotropic effects of mutations on traits (with cases spanning from one trait, to two traits with and without pleiotropy, and with different strengths in selection on each trait, **Figure 1D).** Thus, the simulations spanned scenarios from those that are commonly simulated in the population genetics literature (one oligogenic trait adapting to an environmental cline) to previously unexplored scenarios (two traits with pleiotropy).

**Figure 1.**
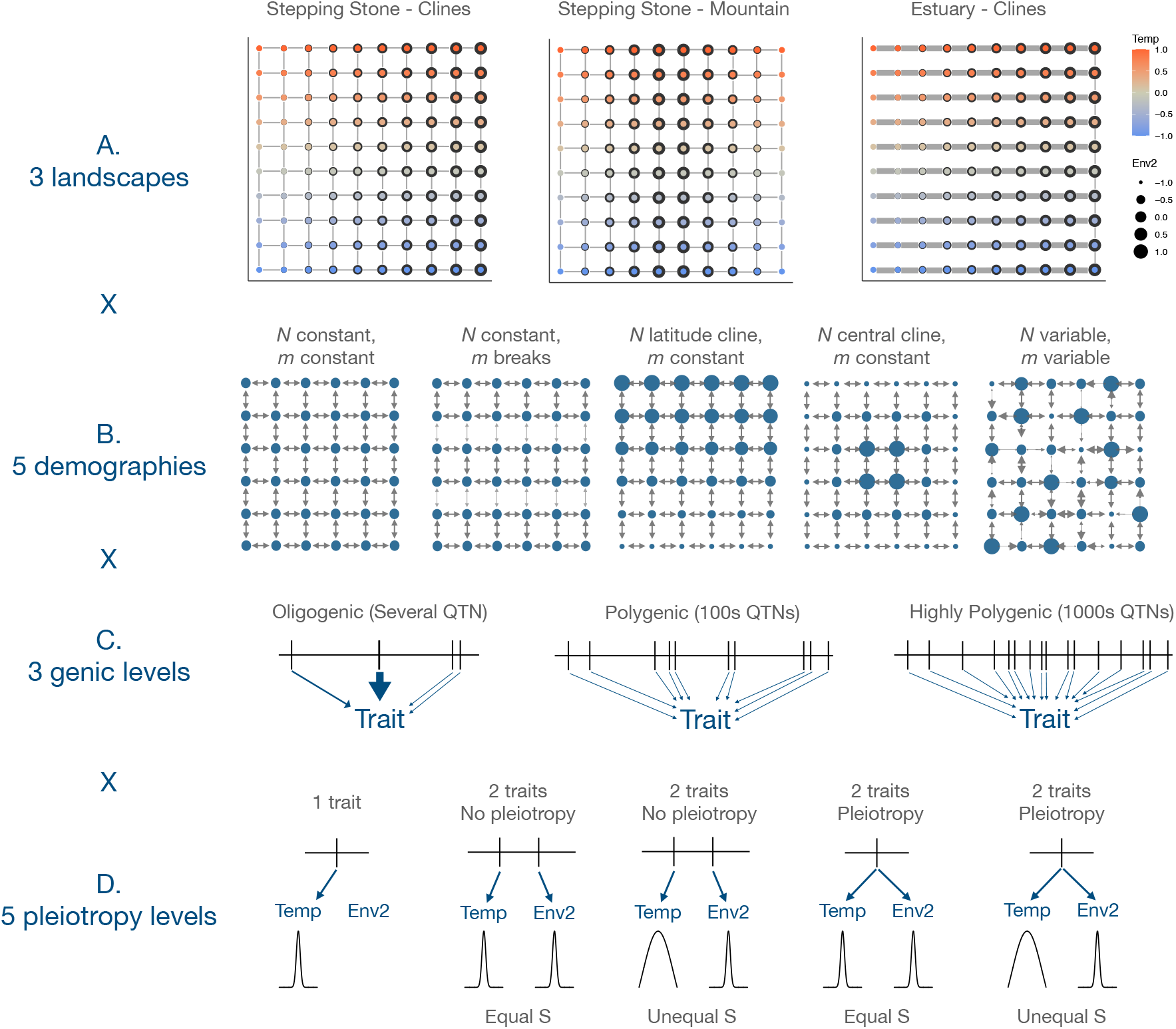
An overview of the parameter levels for the simulations with two quantitative traits. **A)** Three landscapes determined the relationship between patterns of migration and the selective environment. The color of the point indicates the optimum multivariate trait value at that location on the landscape. The optimum “temperature” trait followed a latitudinal cline in all landscapes, while the optimum “Env2” depended on the landscape. Grey lines indicate pathways of migration. **B)** Five demographies determined how drift and migration operated across the landscape (examples show for stepping stone). **C)** Three genic levels determined the number of loci and their effect sizes on the trait. **D)** Five pleiotropy levels determined the number of traits, pleiotropic effects of mutations, and strength of selection on each trait.

The two traits included a “temperature” trait that adapted to a latitudinal environment (in all simulations), and an “Env2” trait that adapted to a longitudinal environment (only in 2-trait simulations). The biological analogy to *Env2* depended on the context: it is analogous to elevation in the *Stepping Stone Mountain* landscape (e.g., trees) or salinity in the *Estuary Clines* landscape (e.g. sticklebacks or oysters) **(Figure 1A).**

The simulations reached an equilibrium level of local adaptation **(Supp. Figure S1),** and clines between each quantitative trait and the selective environment evolved (with correlations ranging between 0.5 and 0.9, x-axis in **Figure 2A).** The amount of divergence was primarily determined by landscape and demography, while the degree of local adaptation (LA) was primarily determined by landscape and genetic architecture **(Supp. Figures S1, S2, S3, S4, Supp. Tables S1, S2).** Across the vast majority of simulations, the population structure (measured as PC1 of the genotype matrix) was highly correlated with deme temperature in all landscapes, but rarely correlated with *Env2* in the *Stepping-Stone-Mountain* or *Estuary-Clines* landscapes **(Supp. Figure S5).** Oligogenic, moderately polygenic, and highly polygenic architectures evolved on average 12, 646, and 3042 causal loci, but this was reduced to 8, 58, and 499, respectively, after MAF filtering **(Supp. Figure S6).** Additional information about the simulations can be found in the Supplemental Results.

**Figure 2.**
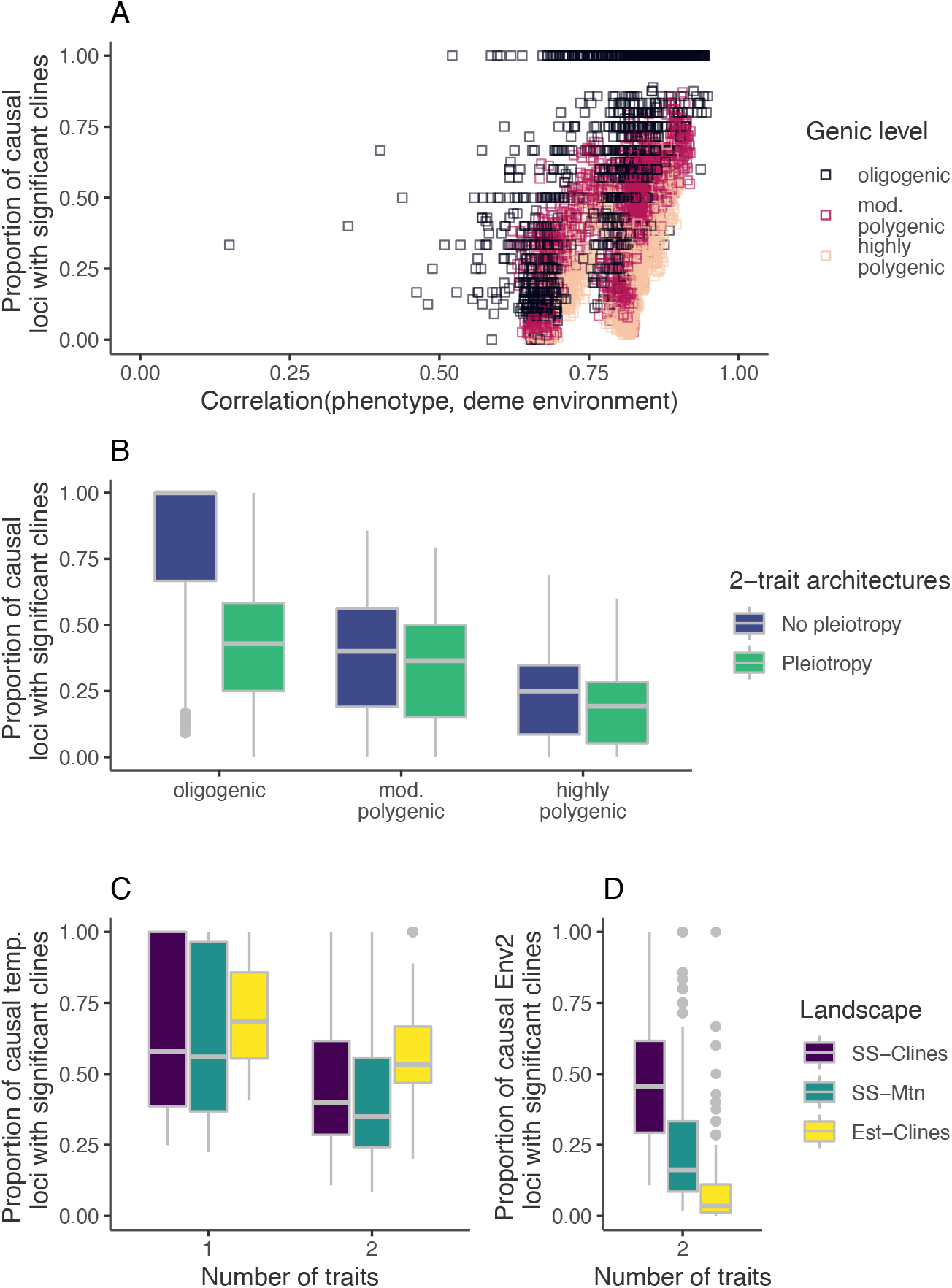
Effect of different parameter levels on the proportion of causal loci that evolved significant clines. **A)** Scatterplot of the proportion of causal loci that have significant allele frequency clines as a function of the degree of the evolved cline in the trait under selection. **B)** The effect of genic level and pleiotropy level, averaged over both traits. **C)** The proportion of causal loci that evolved significant clines in the temperature trait, split out by landscape (color) or by whether the trait was simulated alone or in combination with a second trait (Number of traits). **D)** The effect of the landscape on the proportion of causal loci that evolved significant clines in the Env2 trait (which was only simulated in combination with the temperature trait). Abbreviations: *SS* – *Clines*: stepping stone landscape with clines in both environments; *SS – Mtn*: stepping stone landscape with clines in temperature environment but mountain range for Env2; *Est* – *Clines*: estuary landscape with clines in both environments. See Figure 1 for landscapes.

### Q1: How do phenotypic clines evolve without clines in causal allele frequencies?

#### Interactions among landscape, demography, and genetic architecture determine the proportion of QTNs with clinal patterns

Despite high correlations evolving between environmental phenotypes and deme environments across all simulations, the simulations that evolved allele frequency clines at a large proportion of causal loci were simulations with oligogenic architectures and no pleiotropy **(Figure 2A, B).** The proportion of causal loci that showed significant allele frequency clines decreased as the genetic architecture became more polygenic **(Figure 2B).** There was an interaction between genic level and pleiotropy, as pleiotropy reduced the proportion of loci that exhibited allele frequency clines by ~40% in oligogenic architectures, but this effect became less pronounced as the architecture became more polygenic **(Figure 2B,** compare bars within genic level). In addition, the proportion of causal alleles that evolved clines decreased as the strength of selection became weaker, but was not substantially affected by demography **(Supp. Figure S7).**

When only the temperature trait was simulated alone, a similar proportion of causal temperature QTNs evolved clines across all landscapes **(Figure 2C,** compare medians within “1 Trait”), which is consistent with the relationship between the temperature trait optimum and spatial location being the same among the landscapes **(Figure 1A).** However, when the temperature trait was simulated in conjunction with the *Env2* trait, a slightly decreased proportion of causal temperature QTNs evolved clines **(Figure 2C,** compare medians for “1 Trait” vs. “2 Traits” in the same color bar).

The landscape greatly affected the proportion of causal *Env2* alleles that evolved clines. In the *Stepping-Stone-Clines* landscape, the proportion of causal *Env2* alleles that evolved clines was similar to the proportion of causal temperature alleles that evolved clines (compare dark purple bar in **Figure 2D to 2C,** “2 traits”), which is consistent with this landscape being symmetrical with regard to gene flow and both environments. However, the proportion of *Env2* alleles with clines decreased in the *Stepping-Stone-Mountain* scenario, and was greatly reduced in the *Estuary-Clines* scenario **(Figure 2D).**

#### Phenotypic clines evolved without clines in the underlying allele frequencies under conditions that promoted unique combinations of mutations to evolve to the multivariate optimum in different parts of the landscape

Consider an example from the *Estuary* demography with a pleiotropic and moderately polygenic architecture: phenotypic clines evolved in both traits, but with more variance for the *Env2* trait that experienced higher gene flow **(Figure 3A,B).** Many temperature QTNs exhibited non-monotonic relationships between allele frequency and deme temperature **(Figure 3C),** while most *Env2* QTNs showed no apparent pattern **(Figure 3D).** These apparent patterns were caused by unique sets of alleles that evolved in response to the unique multivariate optima at each inner estuary site **(Figure 3E, F);** a phenomenon that was exacerbated by lack of gene flow between the inner estuary sites. The same phenomenon was observed in the *Stepping-Stone-Mountain* simulations, except in this case unique genetic architectures evolved to the same multivariate optimum in the upper corners or lower corners of the landscape (e.g convergent evolution for the phenotype), because of lack of gene flow between these locations (49).

**Figure 3.**
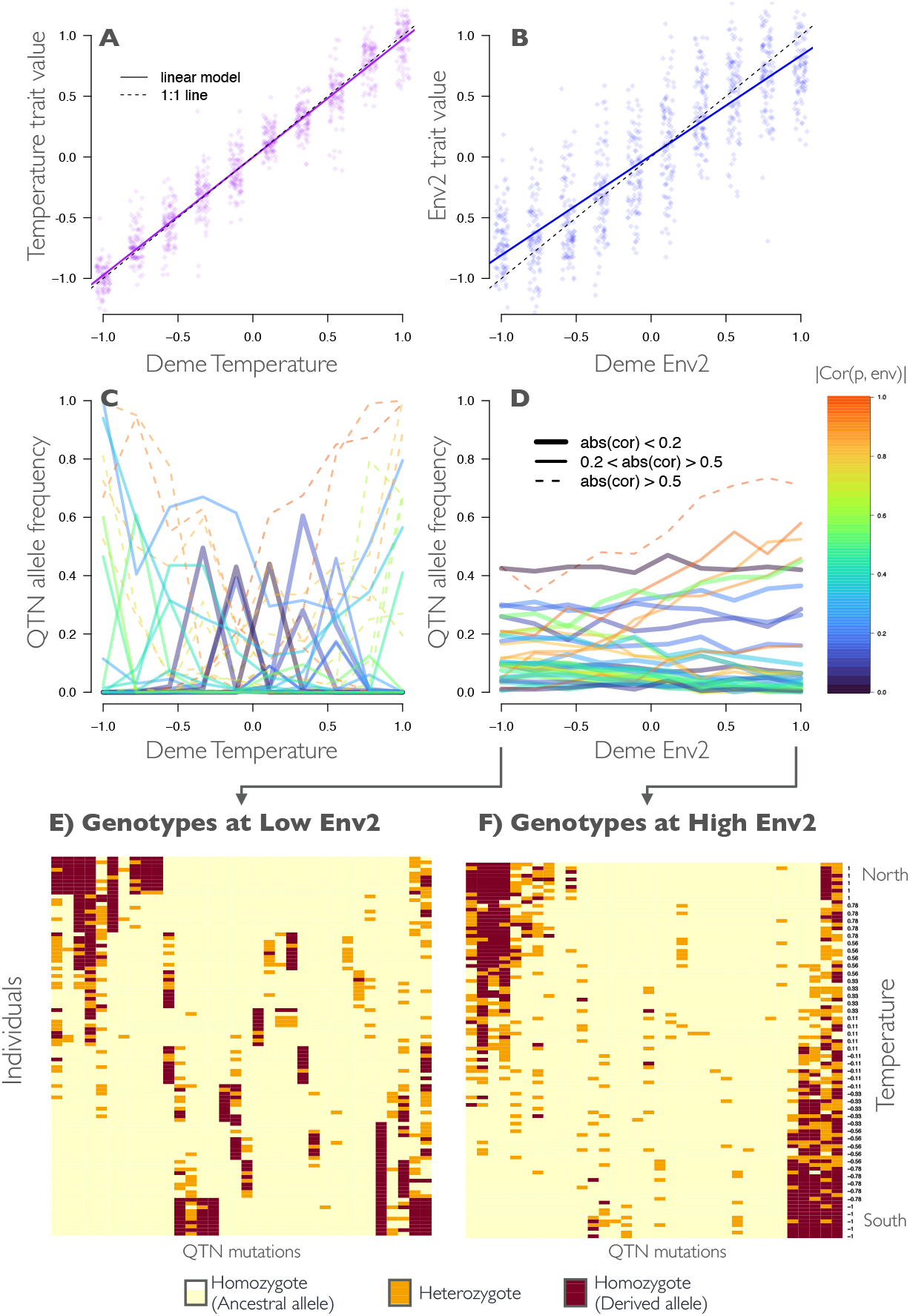
Evolution of phenotypic clines without clines in the underlying allele frequency from a single Estuary-clines simulation. **A) and B)** The evolved individual trait value (i.e., the sum of QTN effect sizes) as a function of deme environment for A) temperature and B) Env 2. **C) and D)** Frequency of the derived allele at each QTN vs. deme environment for C) temperature and D) Env 2. Lines types and colors indicate the strength of the correlation between the derived allele frequency (p) and the environment (|*Cor*(*p, env*)|). **E) and F)** Genotype heatmaps for individuals (in rows) at QTNs (in columns) sampled at all E) low values of Env2 or F) high values of Env2 sites. Individuals ordered from those sampled in the north (high temperatures) to the south (low temperatures). The blocks of derived alleles in (E) shows how specific mutations arose within each inner estuary site that brought the deme closer to the multivariate optimum. Parameter levels: Estuary clines landscape, moderately polygenic, 2 traits with pleiotropy and equal selection strength, *N* central cline, *m* constant, seed 1231214. All QTNs with minor allele frequency > 0.01 are plotted in C-F.

As architectures became more polygenic, a larger number of genetic routes were available to achieve the same phenotype (e.g., high genotypic redundancy, sensu 50). This led to more unique sets of alleles evolving to the multivariate optima in each deme, and hence fewer allele frequency clines. Genotypic redundancy, however, was not a prerequisite for lack of allelic clines. Under oligogenic architectures with low genotypic redundancy, pleiotropy led to fewer allelic clines **(Figure 2B)** because a pleiotropic mutation in a single patch could bring that deme closer to the multivariate optimum in that patch: pleiotropy promoted unique combinations of mutations to evolve to the multivariate optimum in different demes, which reduced GEA patterns.

### Q2: To what extent do GEAs accurately infer the genetic basis of adaptation?

Each simulation was analyzed with commonly used GEA methods: Kendall’sτ rank correlation between genotype and environment (no structure correction), latent factor mixed model [LFMM, univariate, includes structure correction (27,28,32)] and redundancy analysis [RDA, multivariate ordination, with and without structure correction, (36,38,39,51)].

While structure correction greatly reduced false discovery rates for the univariate models (comparing correlation to the LFMM model), it did not reduce false discovery rates for the multivariate RDA **(Figure 4A, Supp. Figure S8).** Power to detect QTNs was generally less than 60% across all approaches, and power was reduced by ~30-50% after structure correction for the temperature model **(Figure 4B).** This occurred because structure correction caused power to decline as the correlation between structure and environment increased (especially at a correlation > 0.25, **Figure 4C, Supp Figure S9),** and temperature was highly correlated with structure across all landscapes **(Supp. Figure S5).**

**Figure 4.**
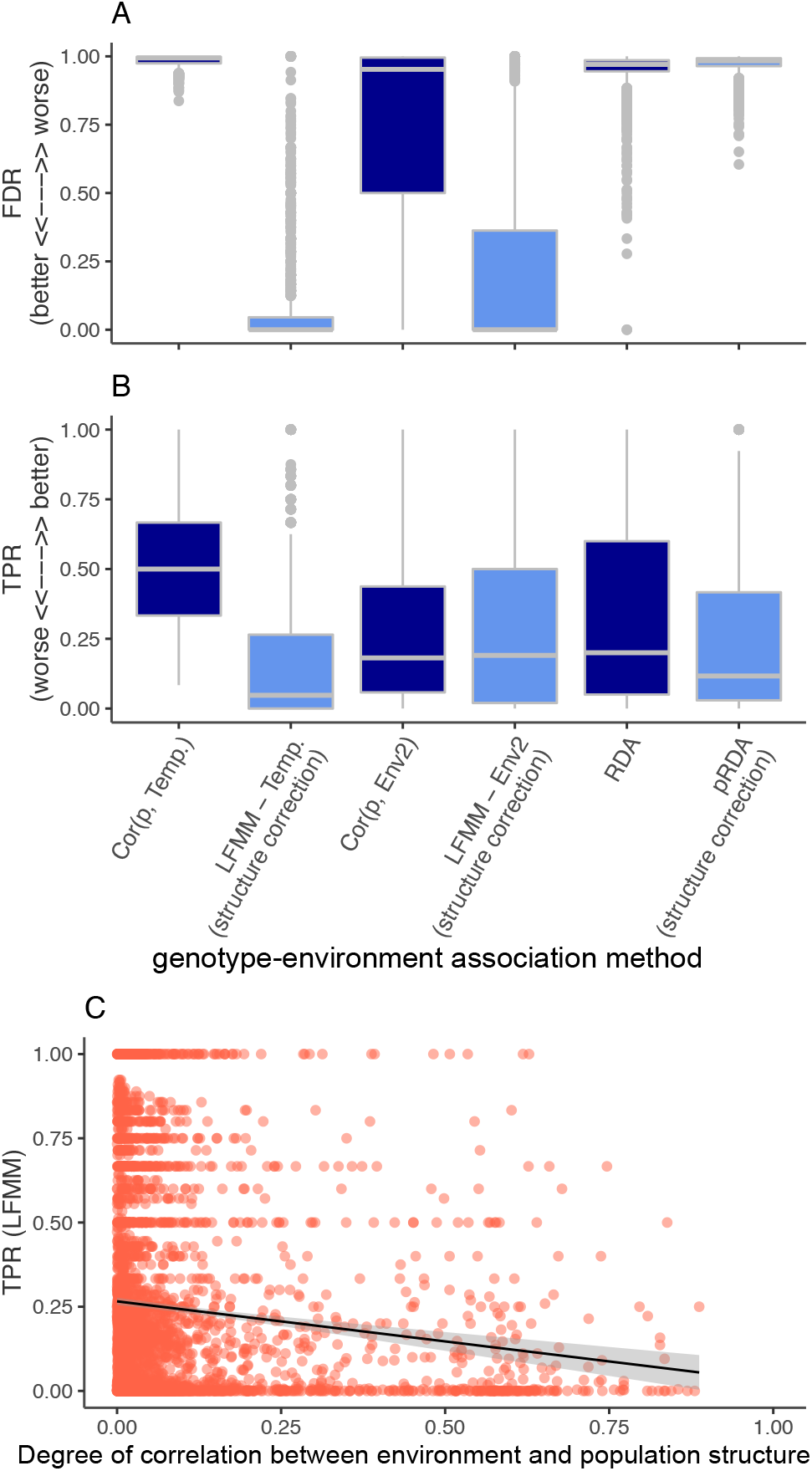
Performance of different methods across all simulations. **A)** False Discovery Rate (FDR) measures the proportion of significant hits that are false positives. **B)** True positive rate (TPR) measures the proportion of QTN loci that are true positives. Note that the temperature environment was highly correlated with structure across all demographies, and the decrease in power after structure correction. **C)** Relationship between the true positive rate of LFMM and the degree to which population structure was correlated with the environment. Population structure was approximated as the first latent factor *U*1 from LFMM. Abbreviations: *Cor*(*p,env*): Correlation between derived allele frequency *p* and the environment; LFMM: latent factor mixed models; (p)RDA: (partial) redundancy analysis.

### Q3 How important are clinal QTNs in adaptation?

Some forecasting models use identified adaptive loci to predict population maladaptation to climate change (52). The heatmaps in Figure 2 suggest that non-clinal QTNs contribute to adaptation in specific demes, so it is possible that clinal QTNs with broadscale geographic patterns are sufficient to explain local adaptation in the metapopulation. Thus, a central question is the extent to which clinal QTNs identified by GEAs make accurate predictions.

I address this question from two perspectives. First, I asked if the proportion of additive genetic variance (*V_A_*) explained by clinal QTNs was different from a null expectation based on the proportion of QTNs that were clinal. Across most of the simulations, the proportion of *V_A_* explained by clinal QTNs was generally higher than the null expectation **(Supp. Figure S10).** This general pattern, however, did not hold for the *Env2* trait in the *Estuary-Clines* landscapes, in which clinal QTNs typically explained less than 40% of the *V_A_* and this was generally not more than the null expectation **(Supp. Figure S10-C).** Although clinal QTNs generally contributed more to *V_A_* than expected, in some simulations their total contribution to *V_A_* was still low (< 50%), such as in highly polygenic architectures or more complex landscapes **(Supp. Figure S10).**

Second, I asked what proportion of the total local adaptation could be explained by different sets of loci given their evolved effect(s) on the trait(s): (i) a combined set of clinal QTNs for both traits (i.e., outlier QTNs without structure correction), and (ii) a combined set of detectable QTNs from the LFMM models in each environment (i.e., outlier QTNs after structure correction). For oligogenic and moderately polygenic simulations, outlier QTNs without structure correction generally explained > 50% of the local adaptation **(Supp. Figure S11).** For these architectures, outlier QTNs after structure correction generally also explained > 50%, although the effects of structure correction were nuanced **(Supp. Figures S11, S12).** For the highly polygenic architectures, clinal QTNs generally explained > 50% of the local adaptation, but they were not outliers after structure correction and this led to outlier QTNs generally explaining less than 30% of the local adaptation **(Supp. Figure S11).**

### Q4 What does multivariate ordination accurately infer about multivariate adaptation?

RDA had high false positive rates and low power regardless of whether structure was included in the model, indicating that it was not appropriate for the detection of QTNs. Interestingly, however, without structure correction individuals visually mapped correctly onto RDA space according to their multivariate phenotypes (see example in **Figure 5A).**

**Figure 5.**
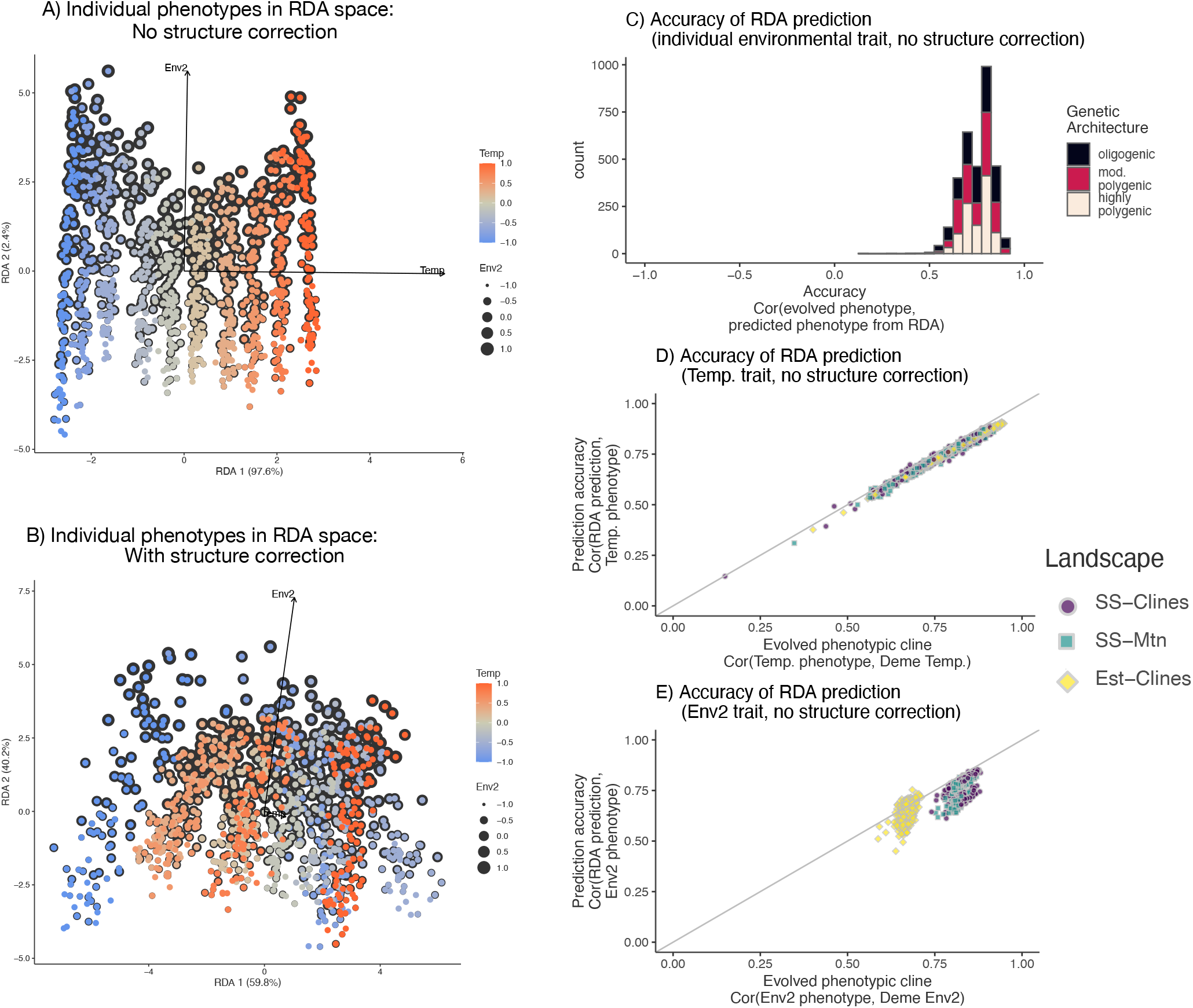
Evaluation of redundancy analysis (RDA) with and without structure correction on the mapping of individuals into RDA space. **A)** RDA. plots with individuals colored according to their evolved trait value (the RDA is based only on genotypes and environment) with (bottom) or without (top) a correction for structure. Parameter levels: Estuary clines landscape, moderately polygenic, 2 traits with pleiotropy and equal selection strength, *N* central cline, *m* constant, seed 1231214. **B)** Across both traits, accuracy of the RDA-predicted individual trait value from Eq. 2 based on that individual’s score in RDA space. Accuracy is measured as the correlation between the evolved trait value and the RDA prediction. **C) and D)** Accuracy of the RDA prediction as a function of the degree that the trait evolved a clinal relationship with the environment for C) Temperature and D) Env2.

I quantified the accuracy of this mapping in the RDA, which was only given information about genotypes and environments. Accuracy was calculated as the Kendall’s correlation between the known trait value *z_ij_* (of individual *i* in quantitative trait *j* that adapted to environment *j*; ground truth value based on the additive effect of causal alleles in that individual’s genome) and the RDA-predicted environmental trait value (based on the score of that individual on the environmental variable RDA space), 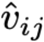:

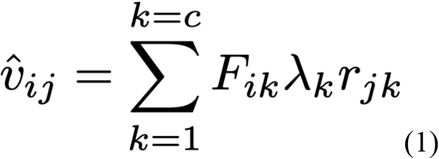

Where 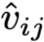 is the RDA-predicted relative environmental trait value (see **Supp. Figure S13** for conceptual visualization), *k* is the canonical axis, *c* is the number of canonical axes (2 in all simulations), *F_ik_* is the “site score” of individual *i* on canonical axis *k*, *λ_k_* is the eigenvalue of axis *k*, and *r_jk_* is the correlation between environment *j* and axis *k* (notation following 53 p. 579-593). The prediction was calculated from an RDA based on 20000 loci randomly chosen from the genome.

For the basic RDA model without structure correction, across all simulations there was a positive correlation between the RDA-predicted environmental trait for each individual (predicted from its scores in RDA space, Eq. 1) and the ground-truth evolved value **(Figure 5C).** Accuracy was not affected by the genic level of the architecture **(Figure 5C, Supp. Figure S14),** and the prediction was accurate despite unique combinations of alleles leading to the similar known trait values. This result occurred because the underlying QTN mutations mapped onto RDA space according to their effect size, and phenotypes were a sum of effects of the causal alleles within each individual genome **(Supplemental Results, Supp. Figure S15).** The accuracy of the RDA-predicted trait value was determined primarily by the degree that the trait evolved to be correlated with the environment **(Figure 5D,E)** and secondarily by the demography **(Supplementary Table S3).** Note, however, that the proportion variance explained by each RDA axis did not necessarily reflect how “important” that axis was. For example, due to the higher gene flow for the Env2 trait in **Figure 5A,** the first RDA axis (on which temperature loaded) explained almost all the variation, even though temperature and *Env2* were weighted equally in the fitness calculation.

The pRDA model with structure correction typically had decreased performance for the multivariate trait prediction, because structure correction jumbled the mapping of individuals in multivariate space (compare **Figure 5A** to **5B**). The mapping of QTNs in multivariate space was also jumbled by structure correction **(Supplemental Results, Supp. Figure S15).** The jumbling occurred primarily along the temperature axis, which was the trait correlated with structure across all demographies **(Figure 5A, Supp. Figure S16).**

An accurate prediction of a quantitative trait for an individual from landscape genetic data could have potentially useful applications, particularly for traits such as temperature tolerance that can be hard to measure in a standardized way (54). This raised the question of how few markers would be necessary to make an accurate prediction of a quantitative trait. Across a wide range of scenarios, accuracy of the RDA trait prediction was similar for 5,000 (~15-20% of the simulated genome) randomly sampled loci from the genome as it was for 20,000 loci (~60-70% of simulated genome), although sampling error was higher in the oligogenic case **(Supp. Figure S17).**

## Discussion

Population genetic models of adaptation to a heterogeneous environment typically assume selection acts directly on the locus (2,5–8). This body of theory predicts that allele frequency at loci under selection will evolve to be more extremely associated with the environment than expected by demography (34), which has led to a large number of statistical methods and empirical studies to seek the genetic basis of environmental adaptation through genotype-environment associations.

Given this body of literature, the evolution of trait clines in response to a selective environment with non-clinal patterns in the underlying allele frequencies initially seems like a paradox. The paradox arises through a quantitative genetic model of selection, under conditions that promote (i) unique combinations of mutations to evolve to the multivariate optimum in different parts of the landscape, and (ii) subtle clines at small-effect mutations that may be only statistically detectable at large sample sizes. Much of the existing literature has focused on the latter issue of subtle allele frequency changes in polygenic adaptation (21,55). By adding spatial complexity, this research demonstrates how genotypic redundancy (i.e., multiple possible genotypes that lead to a similar phenotype (i.e., multiple possible genotypes that lead to a similar phenotype; 50) can interact with reduced gene flow (49) to evolve to non-linear patterns between causal allele frequencies and environments. The unique combinations of mutations are analogous to modular genetic architectures, which have been predicted to evolve in complex environments or when modularity allows adaptation to take place in one trait without undoing the adaptation achieved by another trait (56–59). Such complex possibilities were verbally predicted by Barton in 1979 (8), but have not been elucidated until now. This research also demonstrates, however, that genotypic redundancy is not the only mechanism that can lead to such non-linearities. Oligogenic architectures with low redundancy can also evolve non-clinal patterns when pleiotropic effects allow unique combinations of mutations to achieve spatially varying multivariate optimum.

Recent reviews have concluded that polygenic architectures are common in environmental adaptation (26,60) and that pleiotropy is common in adaptive divergence (61), indicating that this paradox could be common. A way to test for the non-linear patterns predicted by this research is to conduct QTL mapping for quantitative traits, and then examine which QTLs do or do not show clines with the environments that select on the trait. A recent study by Mahoney et al. did exactly this, and discovered some modules of QTL loci that showed patterns of increasing and decreasing allele frequencies across an environmental gradient as predicted here, despite trait clines along the environmental gradient (62). If the patterns predicted by the models herein are common in nature, these results raise questions about the utility of current GEA methods for building a complete picture about the genetic architecture of adaptation to complex environments.

With regards to the question of whether the utility of GEA methods should be re-assessed, it depends on the study question and the nuances of the study system. If the study question seeks to accurately infer the genetic basis of adaptation to the environment, GEA methods will be limited primarily by first principles (i.e., the proportion of selected alleles that evolve clines) and secondarily by statistical issues associated with correction for population structure (either riddled with false positives with structure correction, or decreased power with structure correction if structure correlates with the environment). This raises the possibility that GEA methods will only be useful for accurately inferring the genetic basis of adaptation within a limited parameter space. On the other hand, if the study question seeks to predict the degree of local adaptation in the population, this research showed that GEA outliers that are also true positives can be sufficient for that prediction in oligogenic and moderately polygenic architectures. In most of the parameter space, clinal QTNs made proportionally larger contributions to the additive genetic variance of the trait and to the local adaptation calculation than non-clinal QTNs, because the former evolved broad-scale geographic patterns of adaptation while the latter tended to be highly localized to specific demes.

Despite the limitations of GEA methods, this study highlights the potential use of RDA for predicting relative multivariate quantitative trait values, without needing accurate knowledge of the genetic architecture. Interestingly, the trait prediction was accurate even when two individuals had different genetic architectures that gave the same trait value. This occurred because individual alleles mapped onto RDA space according to their multivariate effect size on the traits, and individual trait values were an additive combination of the alleles they possessed. The multivariate trait prediction was more accurate without a correction for structure in the RDA, because structure correction jumbled the multivariate mapping of loci and individuals along the environment that was more correlated with structure.

Biodiversity management in the face of rapid, multivariate climate change in the world’s terrestrial and marine systems remains an urgent societal need (63,64). Many scientific challenges remain in our ability to adequately address this need, including those associated with identifying adaptive units (65), implementing restoration and assisted gene flow (66), and forecasting the vulnerability of populations (67–69). Various methods for genomic forecasting and genomic offset have been proposed to meet these challenges, but many of them incorporate GEA methods to quantify linear relationships between population-level allele frequencies and environments (52,69,70). Results from this study suggest that GEA methods may be sufficient to forecast broad scale geographic patterns, but may be insufficient to make a forecast about specific individuals or populations. Furthermore, gene-targeted conservation approaches have been criticized because of the difficulty in knowing whether the genomic basis of adaptation has been accurately inferred (71–73).

The results herein raise the possibility that using ordination to predict multivariate trait values at the level of the individual could be a more fruitful approach than GEA methods. Analogous to genomic or marker-assisted selection based on genotype and phenotype (74–77), the multivariate RDA trait prediction based on genotype and environment does not require accurate knowledge of the genomic basis of adaptation, only that linked loci are included in the data. Note that the RDA-multivariate-trait prediction is an unsealed relative value for comparison among individuals, because SNP data only carries relative information between the two alleles at the locus (70). Whether this relative RDA-multivariate-trait prediction will be accurate in more complex environments, in the presence of nuisance variables such as unselective environments, with dominance, and/or with trait plasticity remains an important direction for future research. On a positive note, RDA has been shown to be robust to recombination variation (78). Before a multivariate trait prediction is used in any kind of forecasting or conservation plan it should be rigorously validated with experiments (reviewed in 79). An important next step for empirical research will be to validate the RDA-multivariate-trait prediction by comparing it to a ground-truth value obtained through experimental measures of the performance of individuals in different environments. If multivariate trait predictions meet an acceptable level of performance through this validation process (reviewed in 79), these predictions could prove useful for genomic forecasting, as well as choosing individuals for restoration or assisted gene flow efforts.

## Methods

### Landscapes and demographies

All simulations consisted of 100 demes arranged on a 10 x 10 landscape grid **(Figure 1A).** The 15 levels of landscape-demography were broadly divided into three landscape categories **(Figure 1A),** each with latitudinal and longitudinal selective environments. The first landscape was a stepping stone landscape with a latitudinal and longitudinal cline (*Stepping-Stone Clines*), which is the most commonly simulated scenario in testing methods (16,27,29–31,34–37). The second landscape was a stepping stone landscape with one latitudinal cline and one non-linear longitudinal mountain range *(Stepping-Stone Mountain*). In this scenario, the northeast and northwest corners of the landscape have the same environment (same for southeast/southwest), and so isolation-by-distance did not correspond to isolation-by-environment. This leaves the potential for unique architectures to arise to the same selective pressure at different geographic locations on the landscape. The third landscape was an estuary landscape with a latitudinal and longitudinal cline (*Estuary Clines*, **Figure 1A),** in which geographic distance did not correspond to genetic or environmental distance. The *Estuary Clines* was designed to give insight into adaptation under high gene flow and repeated independent bouts of adaptation, which is analogous to oysters or sticklebacks that repeatedly colonize and adapt to freshwater environments (the within-estuary sites) connected by gene flow in the marine environment (the outer estuary sites) (e.g., (80,81)). For simplicity, I refer to the latitudinal environment as *Temperature* and the longitudinal environment as *Env2*. In summary, *Stepping-Stone Mountain* had a different environmental pattern than *Stepping-Stone Clines* but the same demography, while *Estuary Clines* had the same environmental pattern as *Stepping-Stone Clines* but different demography **(Figure 1A).**

The demographic parameters were chosen such that different landscapes achieved similar levels of neutral genetic differentiation and levels of local adaptation. For the constant migration scenarios in *Stepping-Stone* landscapes, I simulated equal migration between adjacent demes (*m_x_* = *m_y_* = 0.03 unless otherwise stated). For the constant migration scenarios in the *Estuary Clines* landscape, I simulated low migration between the outermost deme in each estuary (*m_y_* = 0.07), with high migration between adjacent demes within estuaries (*m_x_* = 0.49). Despite higher gene flow, *Estuary Clines* evolved higher levels of genetic differentiation than *Stepping-Stone Clines* due to genetic drift among estuaries (see *Results*).

Within each of the three landscapes, 5 demographies were simulated that described the migration rates and effective population sizes on the landscape **(Figure 1B).** The total metapopulation size (*N_metapop_* = 10,000) was controlled for and approximately constant across these five scenarios:

1. *N-constant, m-constant* is the historically most simulated scenario (16,27,29–31,34–37) **(Figure 1B).**
2. *N-constant, m-breaks* introduced a higher degree of confounding between population structure with the temperature optimum by introducing two latitudinal biogeographic breaks with reduced gene flow **(Figure 1B).**
3. *N-latitudinal-cline, m-constant* confounded genetic drift with the optimum temperature trait by introducing an order of magnitude latitudinal cline in *N* (such a situation may be found in the Baltic Sea, in which the population size of several species declines along a strongly selective salinity cline) **(Figure 1B).**
4. *N-central-cline, m-constant* confounded genetic drift with the optimums for both traits by simulating the largest *N* in the range center **(Figure 1B).**
5. Finally, *N-variable, m-variable* dissociated population structure from the adaptive optimums on the landscape **(Figure 1B).**

See **Supp. Tables S4-S5 and Supp. Figures S18-S19** for more details.

### Genetic map

The Wright-Fisher simulations were based on a quantitative genetic model and a genetic map similar to that used in a recent study (78). The genome consisted of 20 linkage groups of 50,000 sites (82–86). The scaled recombination rate for the simulations (*N_metapop_ r* = 0.01) gave a resolution of 0.001 cM between proximate bases and a total length of 50 cM for each linkage group. This resolution mimicked a SNP chip, in which SNPs were collected across a large genetic map (78). The population-scaled neutral mutation rate for the simulations was (*N_metapop_ μ*= 0.001). QTNs could evolve on the first 10 linkage groups, while on the second 10 linkage groups only neutral loci could evolve. The second 10 linkage groups were used to avoid the possibility of low performance in genome scans due to selective sweeps on neutral loci linked to QTNs (see *Methods: GEA performance*).

### Genetic Architecture and Stabilizing Selection

#### Mutation

The simulations assumed a quantitative genetic model where quantitative trait nucleotides (QTNs) contributed additively to the optimal phenotype for each individual without dominance. The genetic architectures were broadly divided into 3 genic categories: oligogenic (few loci of large effect on the trait), moderately polygenic (dozens to hundreds of loci with intermediate effects on the trait), and highly polygenic (hundreds to thousands of loci with small effects on the trait, **Figure 1C, Supp. Table S6).** For QTN mutations under 1 trait or 2 traits without pleiotropy, the univariate effect size of a new QTN mutation was drawn from a normal distribution with a mean of 0 and standard deviation **σ**_QTN_, which depended on the genic level **(Supp. Table S6).** For QTN mutations under 2 traits with pleiotropy, the bivariate effect size was drawn from a multivariate normal distribution with a standard deviation of **σ**_QTN_ for both traits and no covariance, which gave flexibility for mutations to evolve with effects on one or both traits. In this manner the distribution of effect sizes and linkage relationships among causal mutations was allowed to evolve.

#### Pleiotropy

Within each genic category were 5 levels of pleiotropy and selection: (i) 1 trait (which adapted to the latitudinal cline, the most commonly simulated scenario), (ii) 2 traits without pleiotropy (a QTN’s effect was restricted to one trait) and equal strengths of selection on both traits, (iii) 2 traits without pleiotropy and with weaker selection on the latitudinal temperature trait, (iv) 2 traits with pleiotropy (QTNs could evolve effects on one or both traits) and equal strengths of selection on both traits, and (v) 2 traits with pleiotropy and with weaker selection on the latitudinal temperature trait **(Figure 1D, Supp. Table S7).**

#### Selection

To simulate local adaptation, the trait was subject to spatially heterogeneous stabilizing selection with the optimum for each location in space given by the environment. For each individual in each generation, the fitness was determined by a Gaussian function given the difference between the individual’s phenotype and the optimum at that location. For two traits, the fitness for individual *i* at location {*x,y*} was:

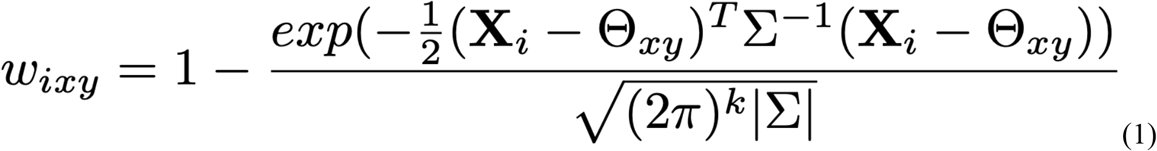

where **X**_*i*_ is a vector of phenotypic values for individual *i* in deme *xy* and **Θ_xy_** is a vector of phenotypic optimums for that deme (optimums shown in **Figure 1A).** Note that equation (1) equals 1 when the individual’s multivariate phenotype is equal to the multivariate optimum phenotype in that patch, and declines away from 1 as the phenotype-optimum mismatch increases. Σ is the symmetric variance-covariance matrix representing the strength of selection on each trait within a deme. For two traits, Σ is a 2×2 matrix with the strength of selection on the diagonals (**σ**_K_ = 0.5 for strong stabilizing selection cases and **σ**_K_ = 4.0 for weak stabilizing selection cases, **Figure 1D, Supp. Table S7)** and zero covariance. For one trait, this equation reduces to the normal distribution.

### Burnin, adding neutral loci, filtering, and sampling

To allow some standing genetic variation at the QTNs to evolve prior to the onset of heterogeneous selection, for the first 1000 generations (bumin), the phenotype was under weak stabilizing selection with a mean multivariate trait optimum (Θ = [0,0]) and weak selection (**σ**_K_ = 4.0) for both traits in all demes. At generation 1000, the environmental optimum at each location changed from the burnin optimum to the final optimum linearly for another 1000 generations, and then remained at the new optimum for the remainder of the simulation (another 1000 generations) during which the phenotype(s) adapted to the optimums in each deme.

Tree sequencing was implemented to record the geneology, and recaptitation with pyslim (v0.6) and msprime (v1.0.1) was used to reconstruct the ancestry of the initial genomes using the coalescent and to add neutral mutations (48,87). VCFTools (0.1.16) (88) was used to filter mutations with a minor allele frequency (MAF) less than 0.01. Then, 10 individuals were sampled from each deme with a probability proportional to their fitness in that patch (to mimic viability selection prior to sampling), for a total sample size of 1000 individuals, and mutations were re-filtered to a MAF < 0.01 based on the sample.

### Quantifying the degree of local adaptation, divergence, and structure

For each replicate, the degree of local adaptation was measured as (i) the difference between population fitness in sympatry and allopatry following (89) and (ii) the correlation between the phenotype and environmental cline for each trait. Overall divergence (genetic differentiation) was calculated as Weir and Cockerham’s *F_ST_* (90) in OutFLANK (91). Population structure was estimated with a principal component analysis on the matrix of genotypes with individuals in rows, loci in columns, and the genotype being the number of derived alleles.

### Quantifying phenotypic clines and identifying clinal QTNs

The degree a phenotype cline was measured as Kendall’s **τ** rank correlation coefficient between individual phenotypes and deme environment. The degree of an allele frequency cline was measured as Kendall’s **τ** rank correlation coefficient (92,93) between deme allele frequency and deme environment, with significance being determined after Bonferroni correction. QTNs that were significant by this criteria were deemed “clinal QTNs.” Heatmaps and other visualizations were used to further understand patterns of adaptation.

### Genotype-environment associations

Latent-factor mixed models (LFMM) assess the linear relationship between genotype and environment. LFMM was implemented using the function lfmm2 in the R package LEA v.4.0.3 (27,28,32). This method estimates allele-environment correlations while simultaneously correcting for population structure with latent factors. The LEA package was also used to conduct a principal components analysis on the genotype matrix, which was used to visualize population structure. The number of population clusters used in analysis for each simulation was based on a scree test (94).

Redundancy analysis (RDA) is a multivariate ordination. RDA aligns multivariate genetic variation with multivariate environmental data to identify the environmental gradients that are the most correlated with genetic variation. Previous oligogenic simulations have shown that RDA can identify causal SNPs as outliers in RDA space (36,38,39,78). The RDA was run with individual-level genotype data as the response variable and the two environments as predictor variables, and models were run both without and with correction for population structure. Correction for population structure was implemented as a partial RDA (pRDA), conditional on the first two principal components axes. Both the RDA and pRDA were implemented using the ‘rda’ function in the R package vegan (95), and outliers were called using the method proposed by (38,39). Note that these simulations present a relatively straightforward scenario for RDA, since the temperature and salinity gradients are orthogonal to each other. Quantities used for the RDA prediction were obtained from the ‘biplot’ output of the ‘rda()’ function.

### GEA performance

The performance of the association metrics were summarized with different performance statistics: (i) false discovery rate (FDR, proportion of outliers that are neutral, lower is better), (ii) true positive rate (TPR, proportion of causal loci that are significant outliers, higher is better), and (iii) the area under the precision-recall curve (AUC-PR, higher is better). In order to provide the most optimistic estimate of a method’s performance, the performance statistics were calculated by only including causal loci and/or neutral loci unaffected by selection (i.e., including only neutral loci on linkage groups 11-20; neutral loci on linkage groups 1-10 could potentially have been affected by selection via linkage disequilibrium with a causal locus, and were excluded). The P-values from the LFMM and RDA tests were corrected by false discovery rate using the ‘qvalue’ package in R (96) to an expected false discovery rate of *q* = 0.05. Note that the expected FDR value is 0.05 given the *q*-value cutoff, but that higher or lower rates occur in practice if the *P*-value distribution is not appropriately calibrated for population structure.

### Importance of clinal QTNs to local adaptation

The proportion of additive genetic variance (*V_A_*) for each QTN was calculated as the additive genetic variance for the focal QTN standardized by the total additive genetic variance following (78). The proportion of additive genetic variance (*V_A_*) explained by clinal QTNs was compared to a null expectation equal to the proportion of QTNs that were clinal.

I estimated the proportion of local explained by different subsets of QTNs: (i) QTNs with MAF >0.01, (ii) clinal QTNs, and (iii) clinal QTNs inferred from latent factor mixed models that include a structure correction. For (ii) and (iii), a GEA model was performed for each environment, and then outlier QTNs were combined into a focal QTN set that was used for the local adaptation prediction. For each focal subset of QTNs, the counts of the derived allele were multiplied by the QTN effect size, summed to get a phenotype, and that phenotype was used in an *in silico* reciprocal transplant to estimate the degree of local adaptation. This estimate was then divided by the total degree of local adaptation (using all QTNs including those below the MAF threshold) to get an estimate of the proportion of local adaptation explained by that focal subset. A proportion near 1 indicates that the focal subset of QTNs explains most of the local adaptation in the data.

## Acknowledgements

Sam Yeaman and Brandon Lind provided helpful discussion and insights that greatly improved the quality of this manuscript.

## Author Contributions

KEL conceptualized the research, wrote the simulation code, ran and analyzed the simulations, and wrote the manuscript.

## Data Accessibility

Upon acceptance of the final manuscript, all data and code will be archived with the Biological and Chemical Oceanography Data Management Office.

## Supplementary Materials

### Supplemental Results

#### Overview of simulations

Correlations between the temperature trait and deme temperature were similar across landscapes, while correlations between the *Env2* trait and deme *Env2* were lower in the *Estuary* scenarios due to higher gene flow along that environment (Supp. Figure S2). All the simulations evolved low-to-moderate levels of divergence with *F_ST_* usually between 0.05 and 0.2 and high levels of local adaptation, with populations having 30-55% higher fitness in sympatry than allopatry (Supp Figure S2, S3, Supp Table S1). Despite the total amount of local adaptation (LA) being slightly higher in the *Stepping Stone* than *Estuary* landscapes (due to higher gene flow in the latter), *Estuary* landscapes had higher divergence due to drift among estuaries (Figure S2, S3). Within a landscape, the degree of local adaptation was determined primarily by the strength of selection on each trait and secondarily by the number of traits under selection (the genic and pleiotropy levels had minimal effects, **Supp. Figure S2).**

The *Stepping Stone* simulations typically evolved 2-3 population clusters, while the *Estuary* simulatios typically evolved 9-10 population clusters (Supp. Figure S4). Most simulations evolved 25,000-35,000 neutral loci after filtering for a minor allele frequency greater than 0.01, except for the *N-variable-m-variable* simulations which had lower diversity (Supp. Figure S5). These simulations with less that 20,000 loci were excluded from the evaluation of the RDA prediction accuracy.

#### Multivariate ordination

Without structure correction the QTN mutations mapped correctly onto RDA space according to their pleiotropic effect sizes (see example in **Supp. Figure S15A, B).** To quantify this, I calculated Kendall’s correlation coefficient between (i) the evolved additive effect on the derived allele at that locus on the quantitative trait (ground truth), and (ii) the unsealed RDA-predicted mutation effect size on the quantitative trait based on the loading of that mutation on the environmental variable in RDA space. This latter measure was calculated as *û_ij_*, the RDA-predicted relative effect size of locus *l* on quantitative traity:

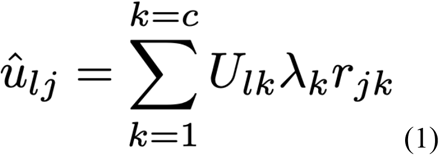

where *k* is the canonical axis, *c* is the total number of canonical axes (2 in all simulations), *U_lk_* is the normalized eigenvector (“species score”) for locus *l* on canonical axis *k*. *λ_k_* is the eigenvalue of axis *k* and *r_jk_* is the correlation between environment *j* and axis *k* (notation following (Legendre & Legendre, 1998)). The measure *û_ij_* is essentially an unsealed back-transformation of the mutational effect from RDA space to the specific quantitative trait *j* that adapts to environment *j* (see **Figure S13-A** for a conceptual visualization).

For the basic RDA model without a structure correction, mutations mapped correctly into ordination space according to their multivariate effects on phenotypes. Across all simulations there was a positive correlation between the mutation effect size predicted from its loadings in RDA space (Eq 1) and the evolved mutation effect size **(Figure 5C).** The accuracy of the prediction decreased as the architecture became more polygenic (e.g. effect sizes became smaller) **(Figure 5C).**

Adding a structure correction to the RDA led to lower accuracy for the RDA-predicted temperature effect size across all conditions **(Figure 5D),** and a slightly higher accuracy for the RDA-predicted *Env2* effect size in the *Stepping-Stone-Mountain* and *Estuary-Clines* landscapes **(Figure 5E).** As a result of the decrease in accuracy for the temperature trait (the trait more correlated with structure), structure correction jumbled how loci mapped into multivariate ordination space, leading to less accurate estimates of multivariate pleiotropic effects of mutations **(Figure 5A).**

Genetic architecture affected the accuracy of the RDA-predicted mutation effect size on the trait, but did not affect the accuracy of the RDA-predicted individual trait value. Although this initially seems like another paradox, the result can be explained by two sources of uncertainty in the trait prediction: (i) the estimated effect size of QTN within the RDA and (ii) sampling error that arose from selecting a subset of markers from the simulation to calculate the trait prediction. The QTN effect sizes for an oligogenic architecture were larger and more accurately estimated, but sampling error increases as the number of QTNs decreases and patterns of linkage around those QTNs become less extensive (as occurred under weaker selection). In contrast, mutation effect sizes for a highly polygenic architecture were smaller and less accurately estimated, but there were hundreds of them and this led to lower sampling error (either through inclusion of a QTN or a neutral locus linked to a QTN) in the multivariate trait prediction. These two sources of uncertainty balanced out in the trait prediction, leading to similar levels of accuracy across architectures for the multivariate trait prediction.

## Supplementary Tables

**Table S1.** The amount of divergence (measured as *F_ST_* and local adaptation (measured as the mean difference of demes in sympatry minus allopatry) for each level of landscape-demography.

| Landscape-Demography | $F_{ST}$ | Amount of local adaptation |
| --- | --- | --- |
| SS-Clines_N-equal_m-constant | 0.06 | 0.48 |
| SS-Mtn_N-equal_m-constant | 0.06 | 0.49 |
| Est-Clines_N-equal_m-constant | 0.09 | 0.43 |
| SS-Clines_N-cline-N-to-S_m-constant | 0.12 | 0.48 |
| SS-Mtn_N-cline-N-to-S_m-constant | 0.13 | 0.48 |
| Est-Clines_N-cline-N-to-S_m-constant | 0.16 | 0.42 |
| SS-Clines_N-cline-center-to-edge_m-constant | 0.13 | 0.48 |
| SS-Mtn_N-cline-center-to-edge_m-constant | 0.14 | 0.48 |
| Est-Clines_N-cline-center-to-edge_m-constant | 0.19 | 0.42 |
| SS-Clines_N-equal_m_breaks | 0.08 | 0.49 |
| SS-Mtn_N-equal_m_breaks | 0.09 | 0.49 |
| Est-Clines_N-equal_m_breaks | 0.13 | 0.43 |
| SS-Clines_N-variable_m-variable | 0.16 | 0.47 |
| SS-Mtn_N-variable_m-variable | 0.17 | 0.47 |
| Est-Clines_N-variable_m-variable | 0.29 | 0.43 |

**Table S2.** The proportion of variance in local adaptation or divergence (measured as *F_ST_*) across all simulations explained by different types of parameters. The proportion of variance was calculated as the sum of squares for each parameter type divided by the total sum of squares in an ANOVA model.

| Parameter type | Proportion of variation in local adaptation explained | Proportion of variation in $F_{ST}$ explained |
| --- | --- | --- |
| Architecture (15 levels) | **0.82** | 0 |
| Demography (5 levels) | 0 | **0.61** |
| Landscape (3 levels: SS-Clines, SS-Mtn, SS-Estuary) | 0.1 | 0.2 |
| Architecture:Demography | 0 | 0 |
| Architecture:Landscape | 0.06 | 0.01 |
| Landscape:Demography | 0 | **0.08** |
| Architecture:Landscape:Demography | 0 | 0.01 |
| Residuals | 0.01 | 0.09 |

**Table S3.** The proportion of variance in different RDA predictions explained by different types of parameters. The proportion of variance was calculated as the sum of squares for each parameter type divided by the total sum of squares in an ANOVA model. The RDA prediction of the mutation effect on the trait is calculated with Eq. 1 in the main paper, the RDA prediction for the trait is calculated with Eq. 2 in the main paper.

| Parameter type | RDA predict:<br>mutation effect<br>on Temp. trait | RDA predict:<br>Temp. trait | RDA predict:<br>mutation effect<br>on Env2 trait | RDA predict:<br>Env2. trait |
| --- | --- | --- | --- | --- |
| Cor(trait, deme environment) | 0.02 | **0.93** | 0.01 | **0.54** |
| Landscape (3 levels) | 0.06 | 0 | **0.1** | 0.03 |
| Demography (5 levels) | 0.01 | 0.02 | 0.05 | **0.22** |
| Architecture (15 levels) | **0.63** | 0 | **0.34** | 0 |
| Landscape:Demography | 0 | 0 | 0.02 | **0.08** |
| Landscape:Architecture | 0.01 | 0 | **0.09** | 0 |
| Demography:Architecture | 0.01 | 0 | 0.02 | 0.01 |
| Landscape:Demography:Architecture | 0.02 | 0 | 0.02 | 0.01 |
| Residuals | 0.24 | 0.03 | 0.34 | 0.11 |

**Table S4.**
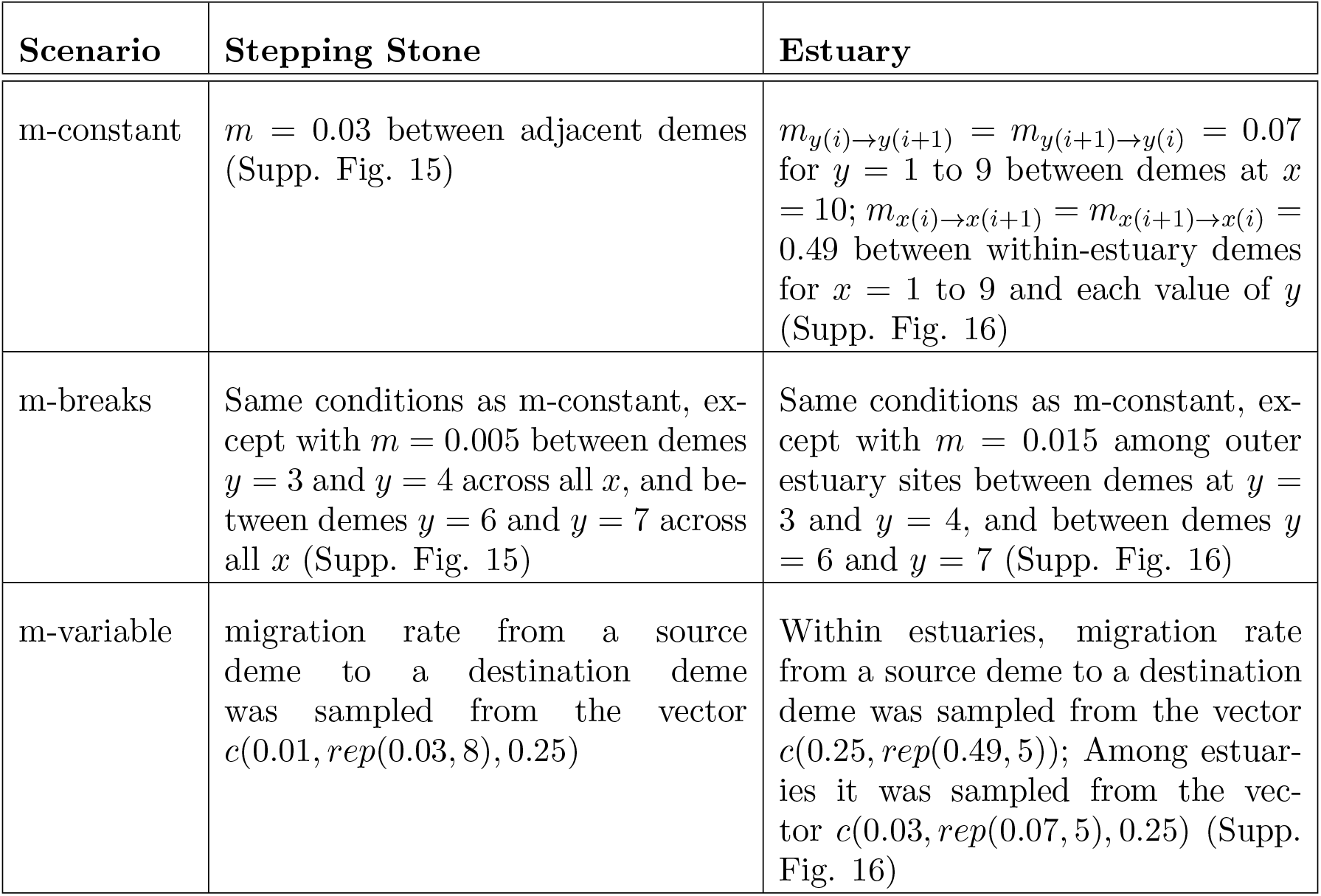
Parameters used for the migration levels. Note that parameters for the Stepping Stone and Estuary simulations were manipulated such that similar levels of neutral genetic differentiation would be generated on the different landscapes. See Supp. Figures 15-16 for visualization.

| Scenario | Stepping Stone | Estuary |
| --- | --- | --- |
| m-constant | $m = 0.03$ between adjacent demes (Supp. Fig. 15) | $m_{y(i) \rightarrow y(i+1)} = m_{y(i+1) \rightarrow y(i)} = 0.07$ for $y = 1$ to $9$ between demes at $x = 10$; $m_{x(i) \rightarrow x(i+1)} = m_{x(i+1) \rightarrow x(i)} = 0.49$ between within-estuary demes for $x = 1$ to $9$ and each value of $y$ (Supp. Fig. 16) |
| m-breaks | Same conditions as m-constant, except with $m = 0.005$ between demes $y = 3$ and $y = 4$ across all $x$, and between demes $y = 6$ and $y = 7$ across all $x$ (Supp. Fig. 15) | Same conditions as m-constant, except with $m = 0.015$ among outer estuary sites between demes at $y = 3$ and $y = 4$, and between demes $y = 6$ and $y = 7$ (Supp. Fig. 16) |
| m-variable | migration rate from a source deme to a destination deme was sampled from the vector $c(0.01, rep(0.03, 8), 0.25)$ | Within estuaries, migration rate from a source deme to a destination deme was sampled from the vector $c(0.25, rep(0.49, 5))$; Among estuaries it was sampled from the vector $c(0.03, rep(0.07, 5), 0.25)$ (Supp. Fig. 16) |

**Table S5.** Parameters used for the population sizes. See Figure 1B for rough visualization.

| Scenario | $N$ on the landscape |
| --- | --- |
| N-constant | $N_{deme} = 100$ |
| N-latitudinal-cline | $N_{deme[y]} = 10, 10, 50, 50, 95, 95, 145, 145, 200, 200$ from $y = 1$ to $y = 10$ |
| N-central-cline | multivariate normal distribution with mean $\mu_x = \mu_y = 4.5$, standard deviation $\sigma_x = \sigma_y = 4.95$, multiplied by $10000 + 5$ |
| N-variable | randomly sampled an $N_{deme}$ value from the vector $10, 10, 50, 50, 95, 95, 145, 145, 200, 200$ |

**Table S6.** Parameters used for the genic levels.

| **Genic level** | $\mu_{QTN}$ | $\sigma_{QTN}$ (same for both traits) |
| --- | --- | --- |
| Oligogenic | $1.0^{-10}$ | 0.4 |
| Moderately Polygenic | $1.0^{-8}$ | 0.1 |
| Highly Polygenic | $2.5^{-8}$ | 0.002 |

**Table S7.**
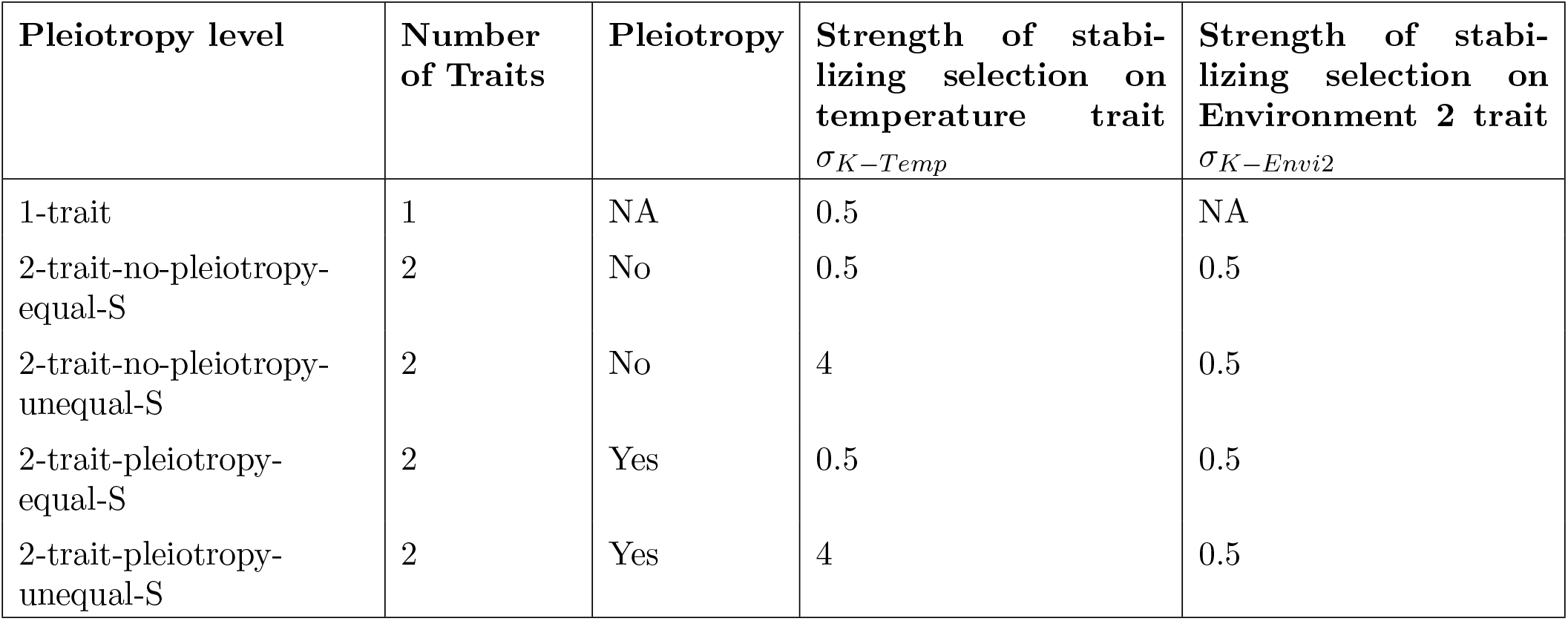
Parameters used for the different levels of pleiotropy and selection strength in the simulations

## Supplementary Figures

**Figure S1.**
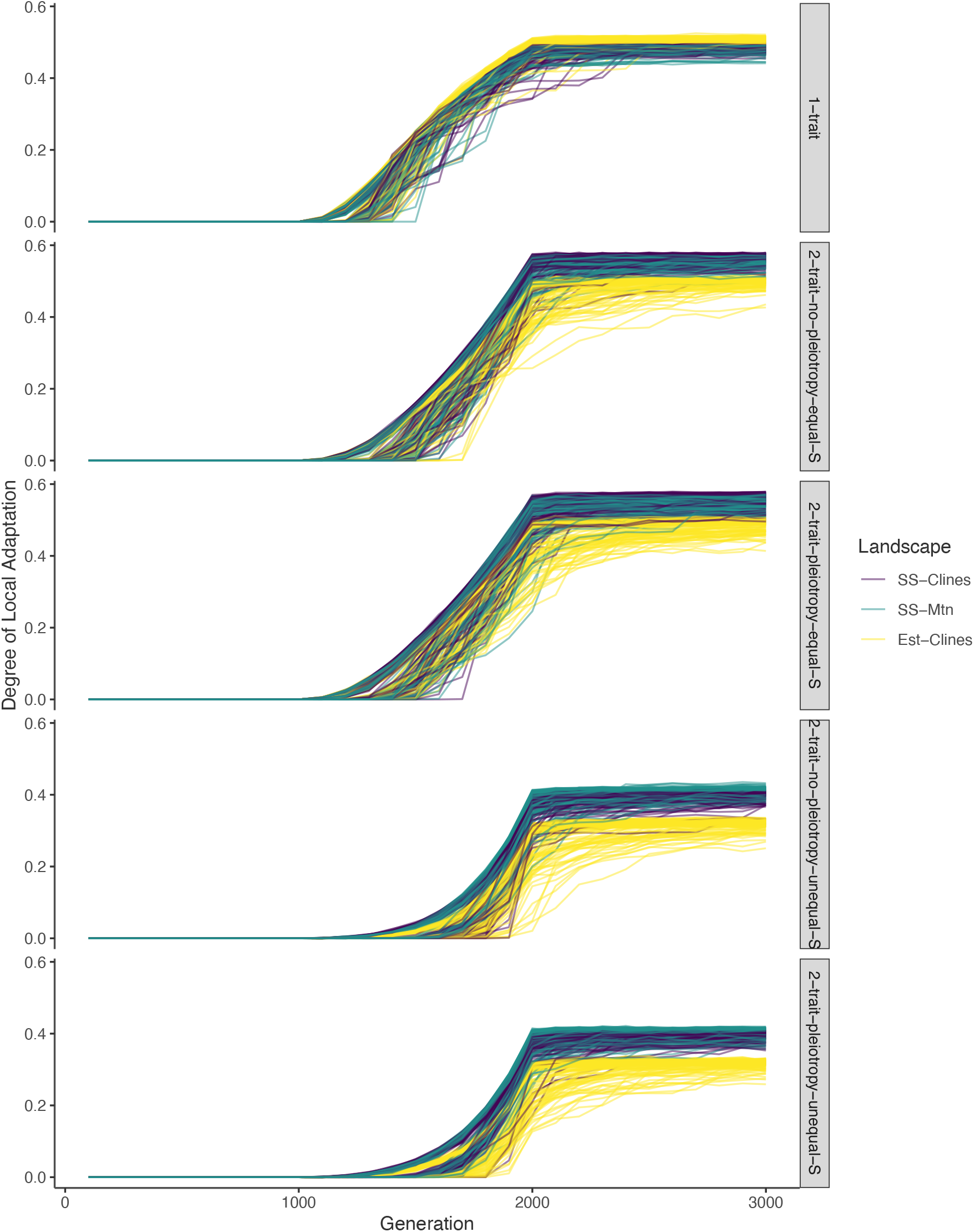
Degree of local adaptation through time, faceted by pleiotropy levels. The first 1000 generations were burnin conditions, in which each deme had the same multivariate trait optimum.

**Figure S2.**
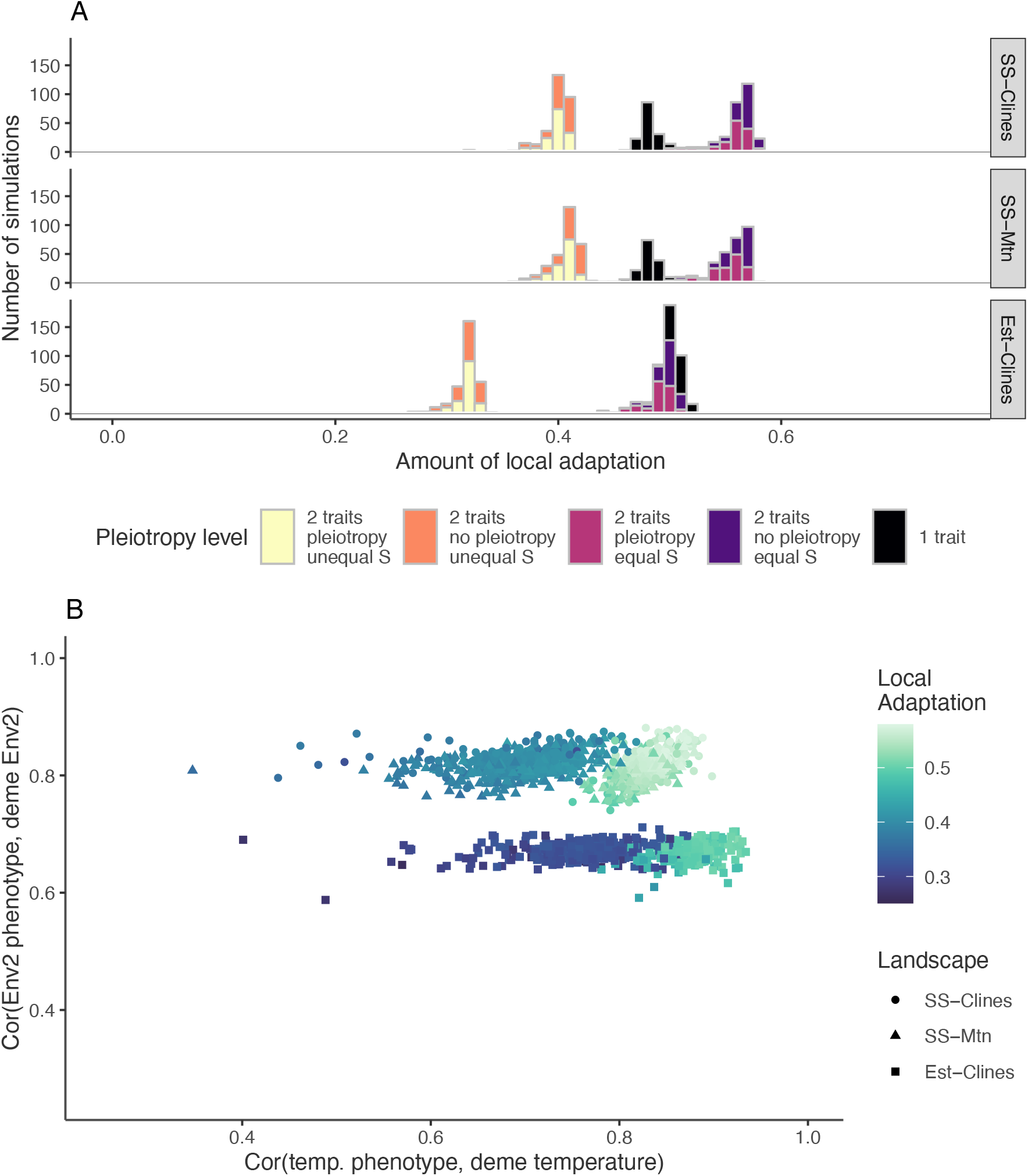
A) A stacked bar plot of the distribution of local adaptation in simulations, faceted by landscape and colored by pleiotropy level. Local adaptation was measured as the mean difference in fitness of demes in sympatry minus their mean fitness in allopatry). B) Distribution of the evolved correlations between the environmental trait and deme environment, for temperature (x axis) and Env2 (y axis). The Estuary-Clines landscape evolved lower correlations for the Env-2 trait because of the higher gene flow within estuaries. Note that in (B) only simulations with both traits under selection are plotted.

**Figure S3.**
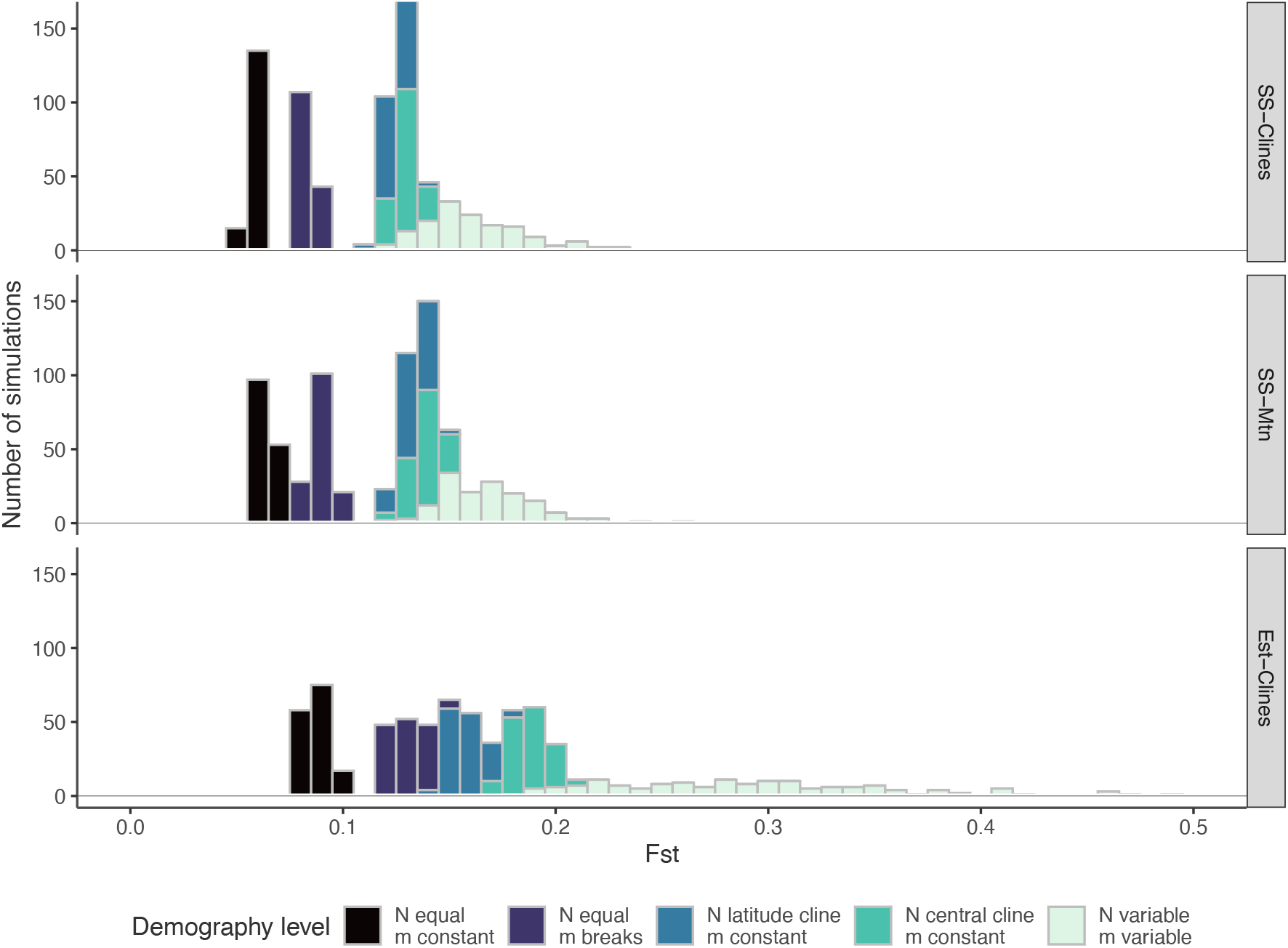
A stacked bar plot of the distribution of divergence in simulations, measured as *F_ST_*, faceted by landscape, and colored by demography.

**Figure S4.**
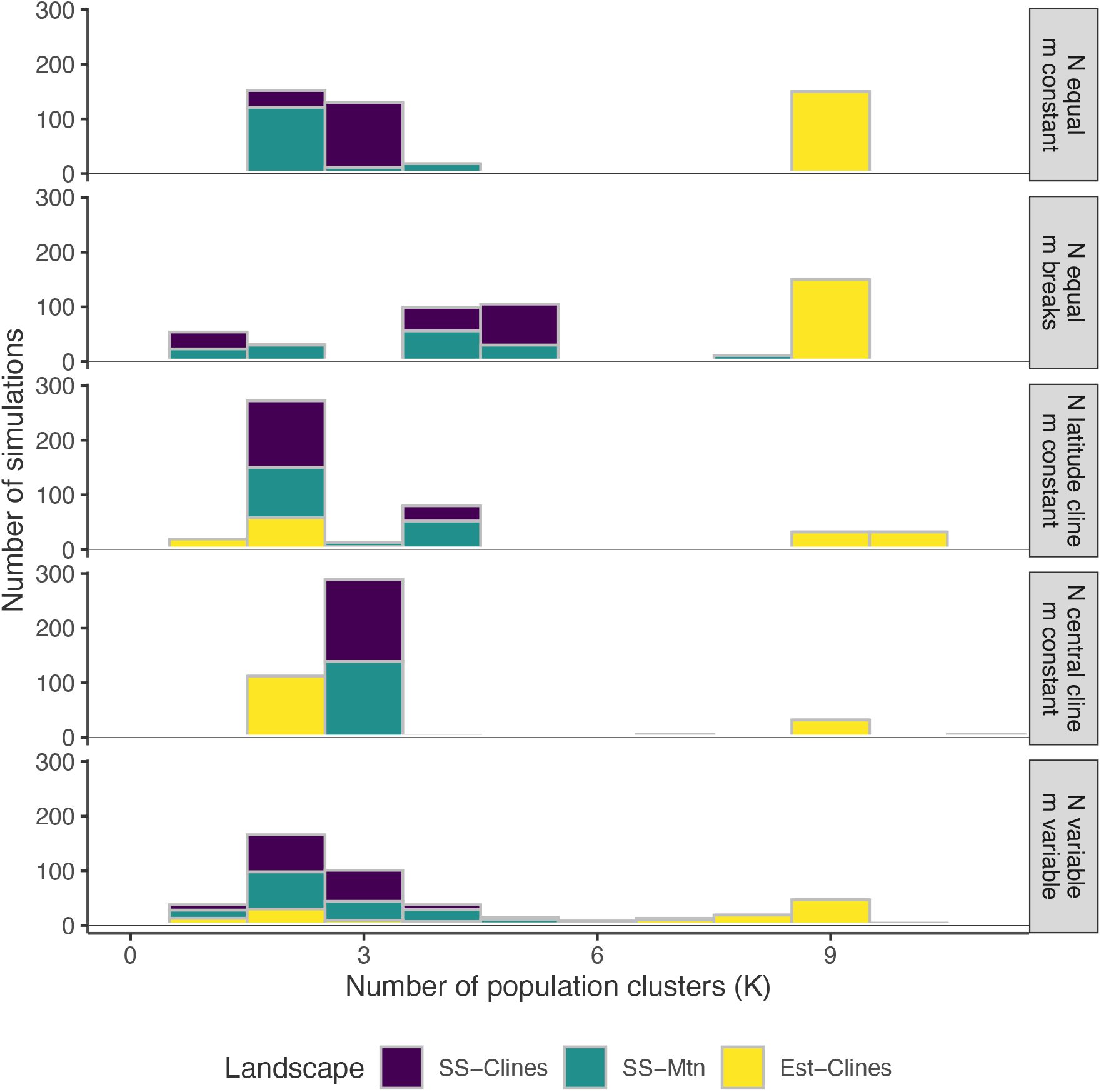
Distribution of the number of population clusters (*K*) in simulations. K was determined by Scree’s rule (see Methods).

**Figure S5.**
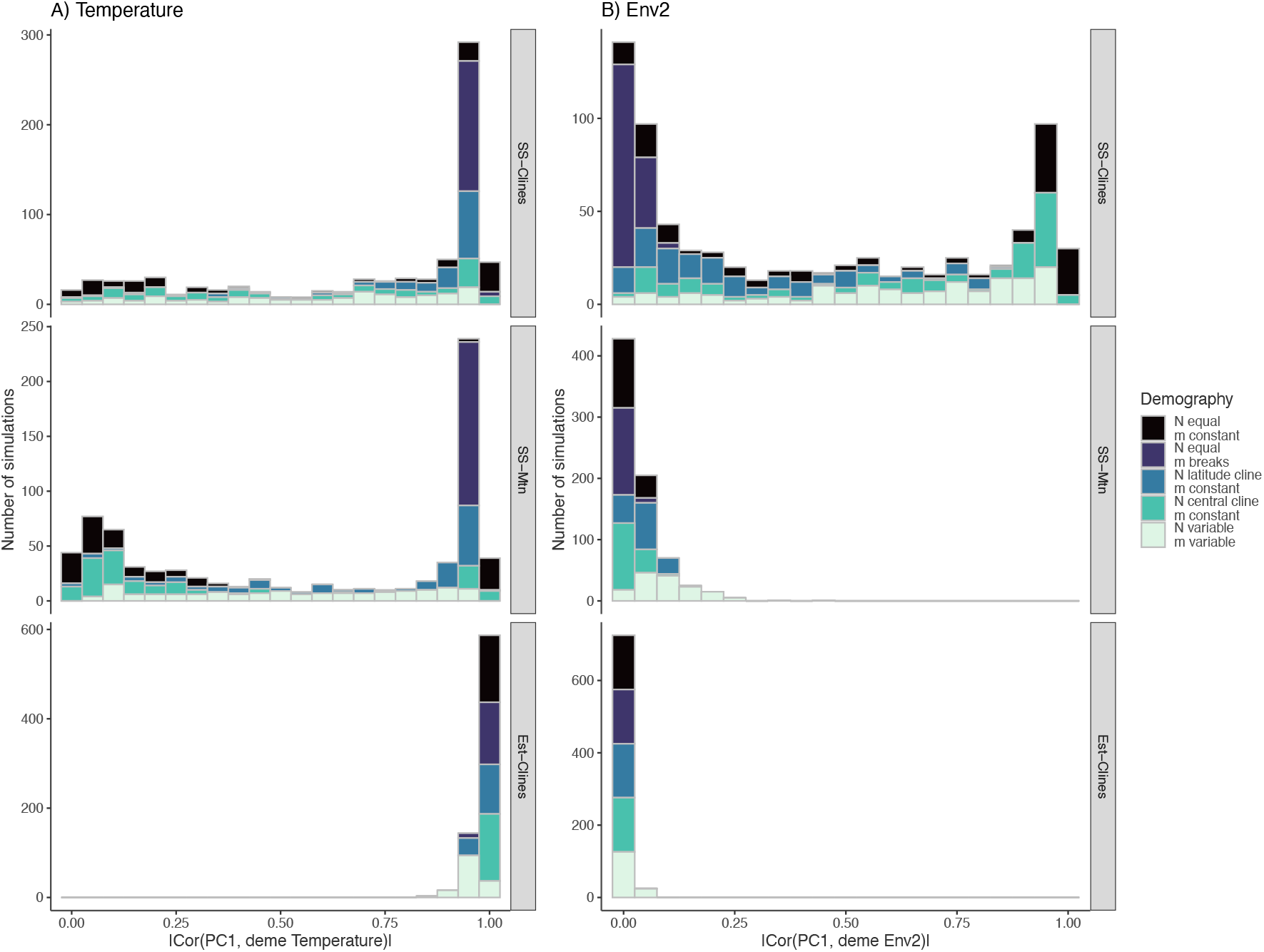
A stacked bar plot showing the distribution of the absolute value of the correlation between population structure and deme environment. Population structure was measured as the first principal component *PC*1 on a principal components analysis of the SNP genotype matrix (with SNPs having a minor allele frequency greater than 0.01).

**Figure S6.**
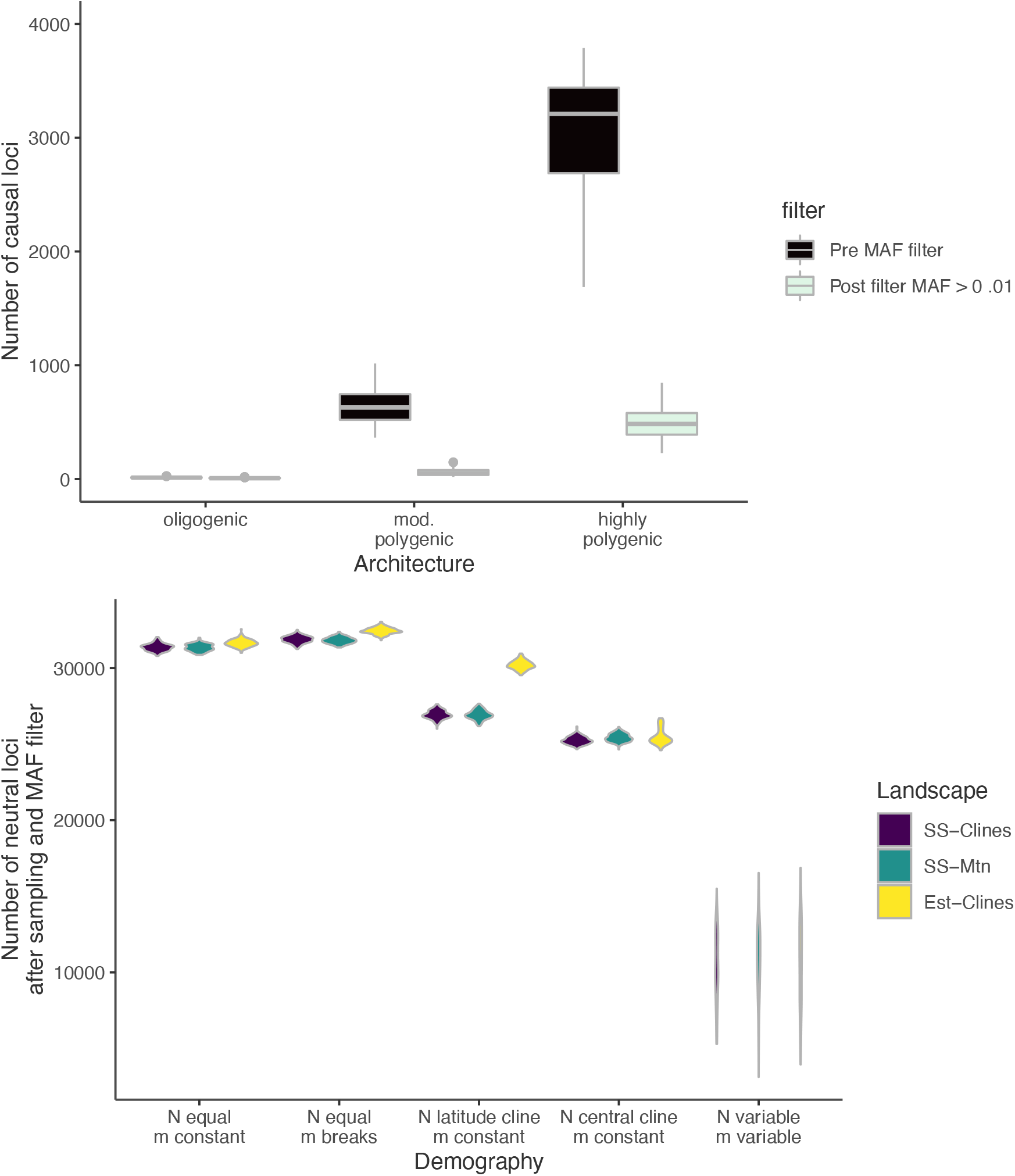
A) Distribution of the number of causal quantitative trait nucleotide loci in the simulations, before and after filtering for MAF < 0.01. B) Distribution of the number of neutral loci in the simulations after MAF filtering < 0.01. The low number in the *N*-variable-*m*-variable simulations reflects the stronger effect of genetic drift relative to other scenarios.

**Figure S7.**
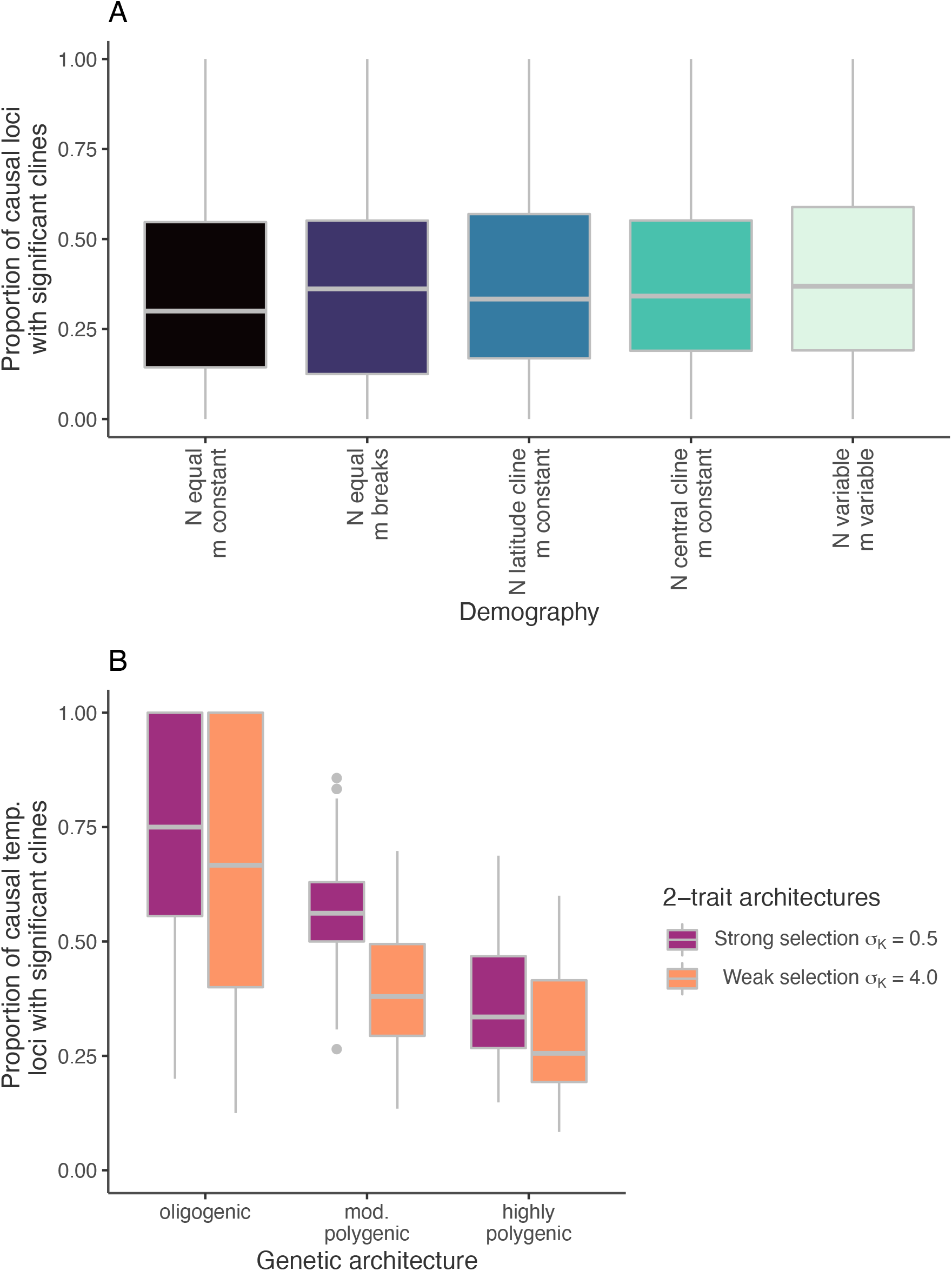
A) Proportion of causal loci with significant clines by demography. B) Proportion of loci with causal effects on the temperature trait with significant clines by the strength of selection on that trait. Only the temperature trait was simulated with strong and weak selection.

**Figure S8.**
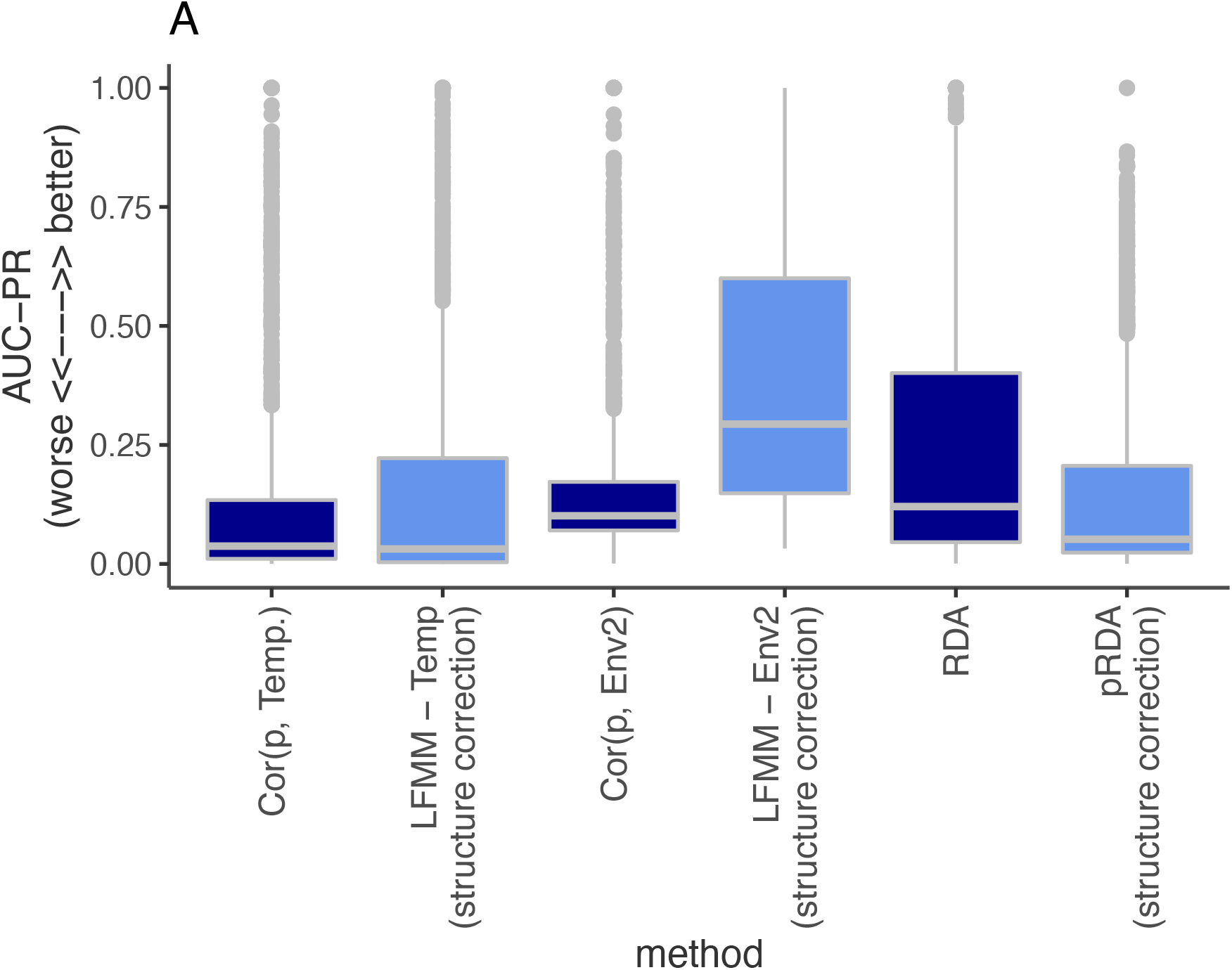
Performance of genotype-environment associations as area under the precision-recall curve (AUC-PR). (AUC-PR) measures the degree to which signals at QTNs are more extreme than neutral loci. A value of 1 indicates all QTN loci had higher signals than neutral loci, a value of 0 indicates the signals at QTN loci rank randomly with neutral loci. This shows how structure correction improved detection of QTNs with effects on the *Env*2 trait. Abbreviations: *Cor*(*p, env*): Correlation between derived allele frequency *p* and the environment; LFMM: latent factor mixed models; (p)RDA: (partial) redundancy analysis.

**Figure S9.**
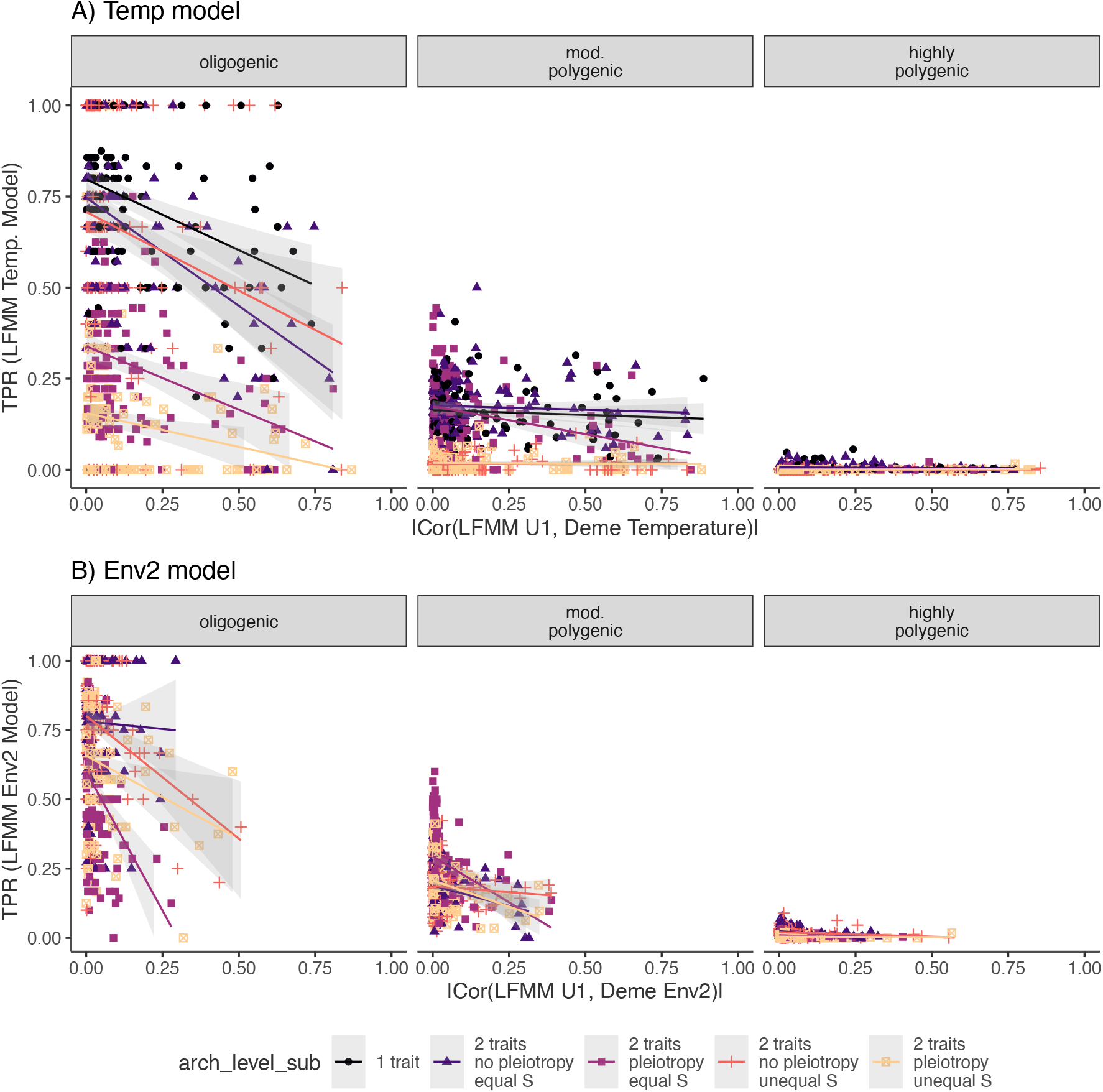
Performance of latent factor mixed model (LFMM) as a function of the how correlated structure was with the environment. This latter quantity was estimated as absolute value of the correlation between (i) the first latent factor of LFMM (U1, i.e. primary axis of structure) and (ii) deme environment. Abbreviations: TPR: True Positive Rate; *S*: strength of stabilizing selection. Performance of LFMM declines as the structure-environment correlation increases because correction for structure lessens the signal of adaptation to the environment.

**Figure S10.**
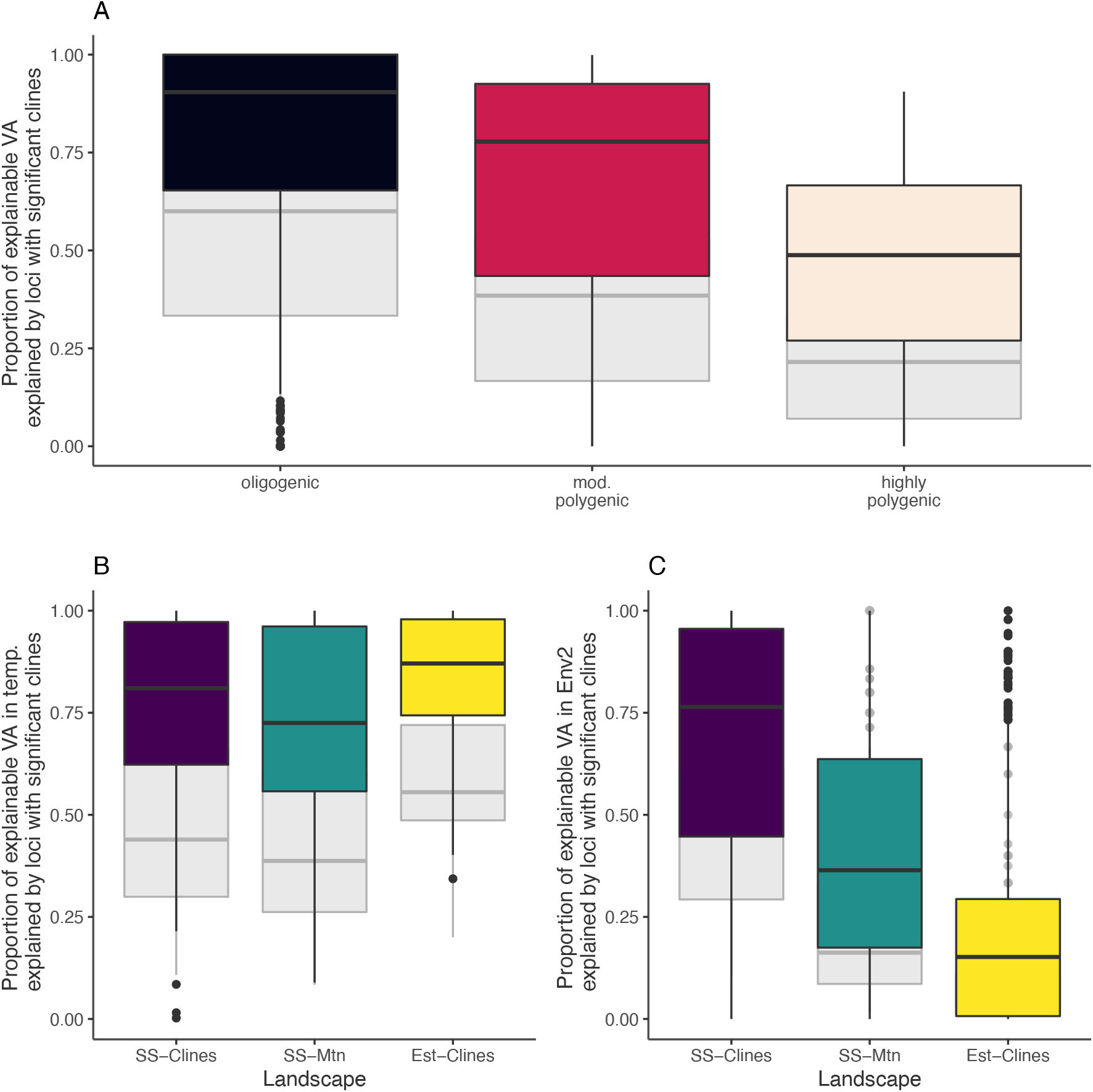
Proportion of additive genetic variance explained by clinal QTNs (colored boxplots), compared to the null expectation based on the proportion of QTNs that are clinal (grey boxplots). (A) The proportion of additive genetic variance (*V_A_*) explained by clinal QTNs as a function of genic level. B) The proportion of additive genetic variance (*V_A_*) explained by QTNs with temperature clines and causal effects on the temperature trait. C) The proportion of additive genetic variance (*V_A_*) explained by QTNs with temperature clines and causal effects on the temperature trait.

**Figure S11.**
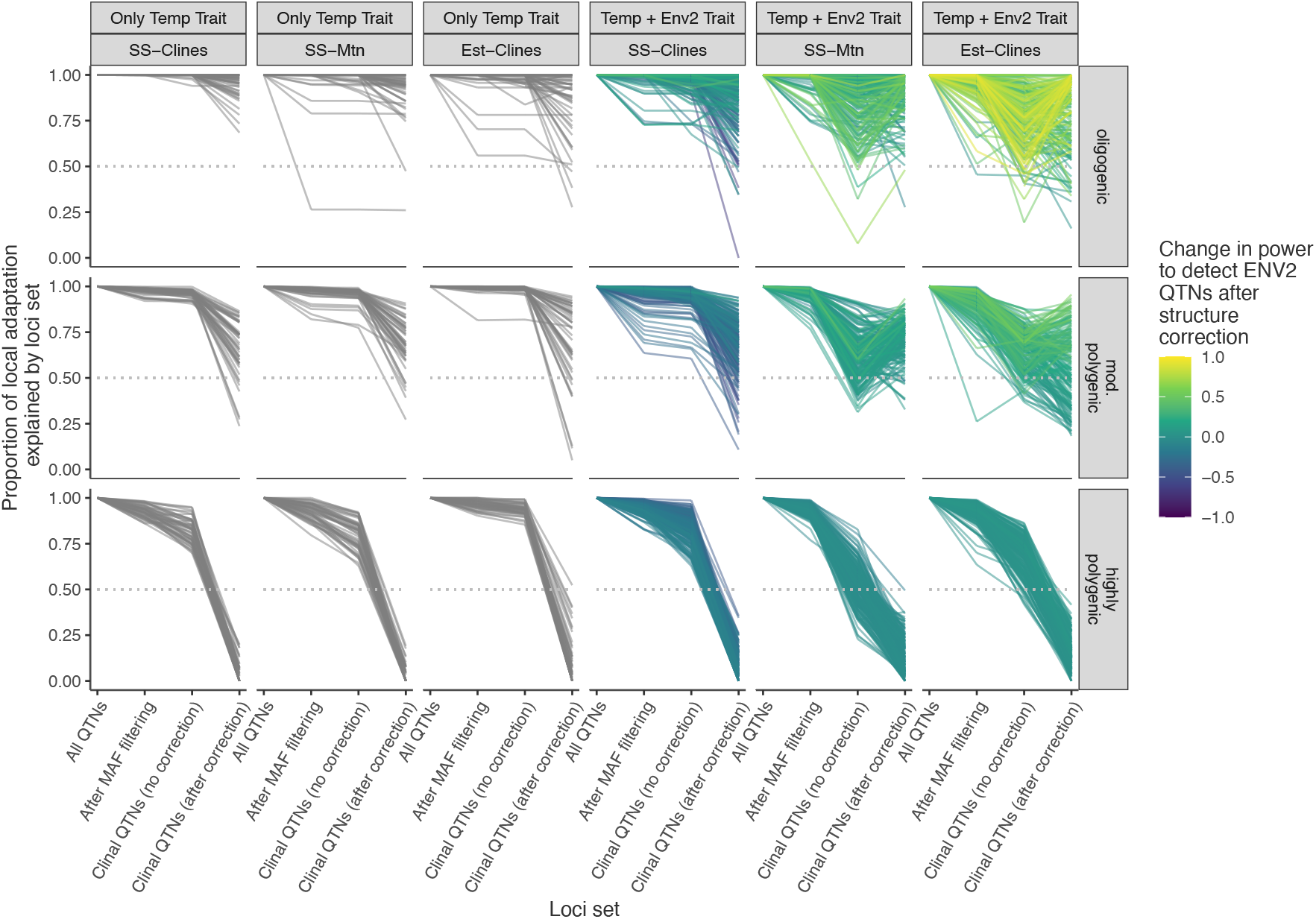
Proportion of local explained by different sets of QTNs (colored boxplots): all QTNs (including those with low minor allele frequency (*MAF*) less than 0.01), QTNs with *MAF* > 0.01, clinal QTNs based on Kendall’s rank correlation between allele frequency and the environment, and clinal QTNs inferred from latent factor mixed models that include a structure correction. Individual GEA models were performed for each environment, and then outlier QTNs were combined into a focal QTN set that was used for the local adaptation prediction. For each focal QTN set, counts of the derived allele were multiplied by the QTN effect size, summed to get a phenotype, and that phenotype was used in an *in silico* reciprocal transplant to estimate the degree of local adaptation. The color ramp represents the change in power to detect Env-2 QTNs after structure correction (i.e., true positive rate for latent factor mixed model between genotype and environment minus the true positive rate for correlation between genotype and environment). In cases where structure correction led to increased power to detect QTNs with effects on the trait adapting to Env2, the local adaptation prediction improved substantially compared to without a structure correction (yellow colors). Abbreviations: *SS* – *Clines*: stepping stone landscape with clines in both environments; *SS* – *Mtn*: stepping stone landscape with clines in temperature environment but mountain range for Env2; *Est* – *Clines*: estuary landscape with clines in both environments.

**Figure S12.**
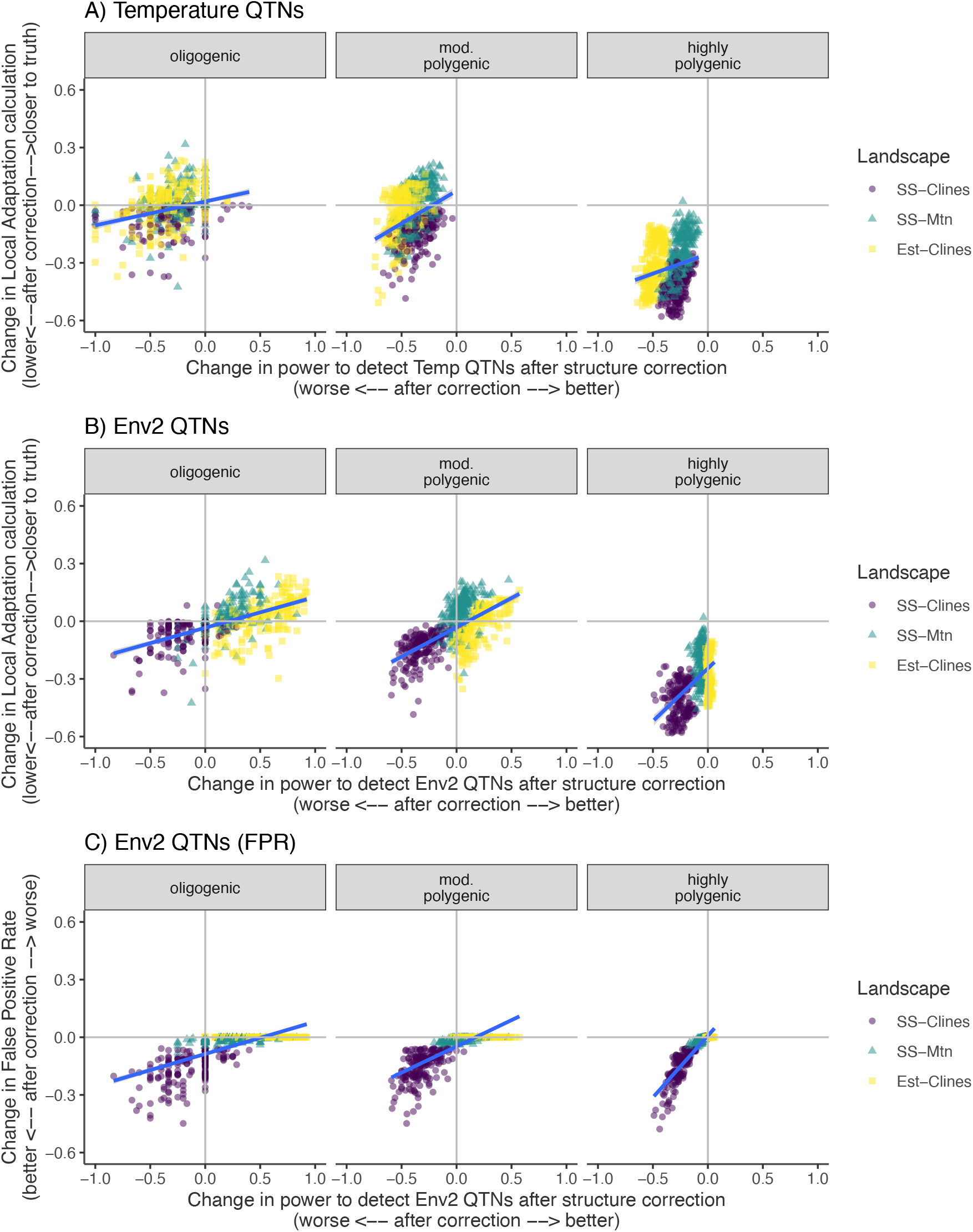
A & B)Understanding how structure correction changes the power to detect QTNs (x-axis) and changes the local adaptation calculation (y-axis). C) When structure correction increases power to detect QTNs for Env2 (x-axis), it does not increase the false positive rate (FPR, y-axis). Abbreviations: *SS* – *Clines*: stepping stone landscape with clines in both environments; *SS* – *Mtn*: stepping stone landscape with clines in temperature environment but mountain range for Env2; *Est* – *Clines*: estuary landscape with clines in both environments.

**Figure S13.**
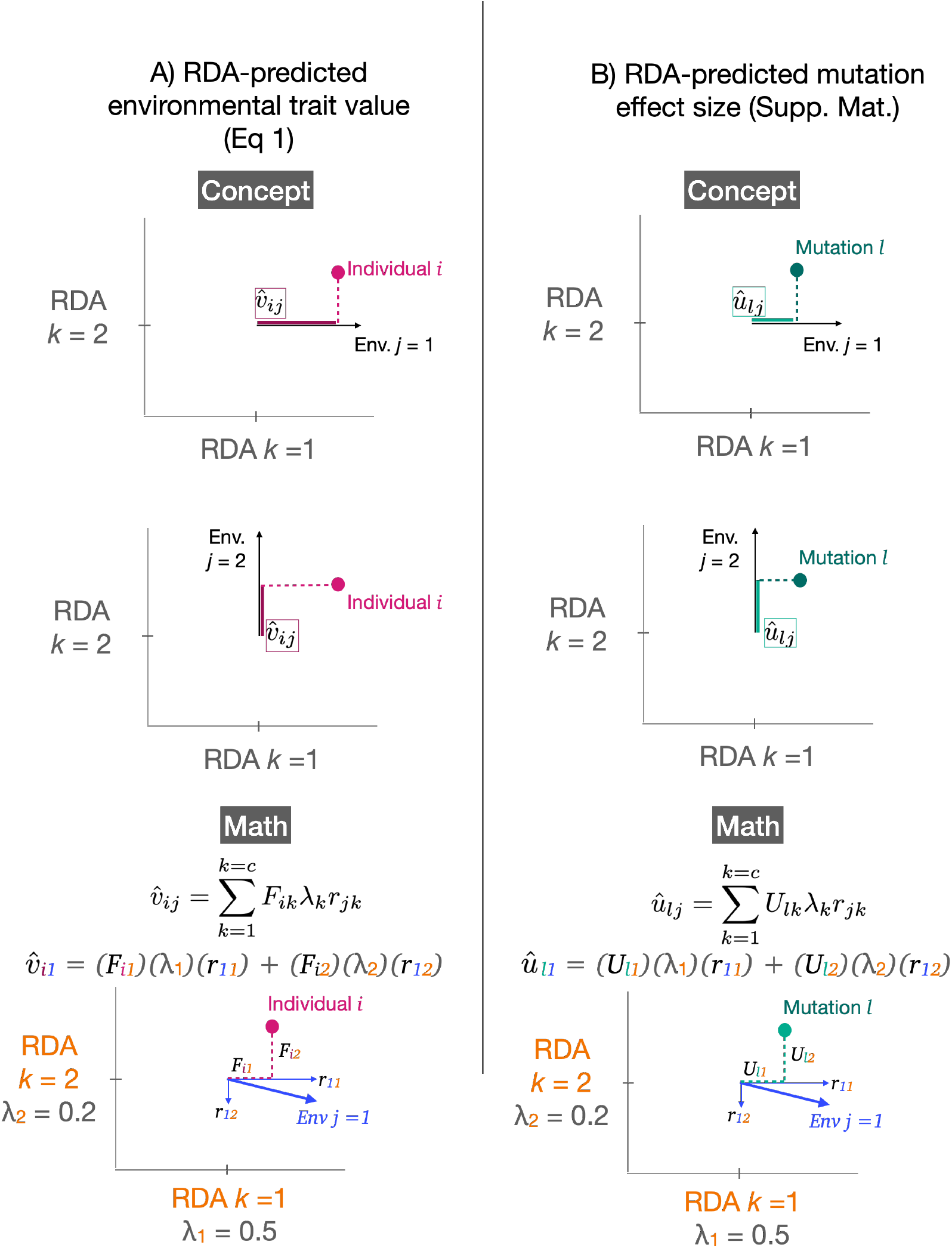
Conceptual visualization of how the mapping of (A) individuals and (B) loci in RDA space can be used to back-calculate the multivariate phenotypes (as in A) or multivariate mutation effect sizes (as in B). A) At the top is a conceptual visualization for the multivariate environmental trait prediction when an environment aligns perfectly with a single RDA axis. In this case the math reduces to the score of that individual on that RDA axis. At the bottom is shown a more complex example when the environment loads onto multiple RDA axes. In this case the environmental trait prediction is a sum of the scores along each RDA axis, scaled by the eigenvalue of each axis and the loading of the environment along each axis. B) At the top is a conceptual visualization for the multivariate environmental mutation effect size prediction when an environment aligns perfectly with a single RDA axis. In this case the math reduces to the score of that mutation on that RDA axis. At the bottom is shown a more complex example when the environment loads onto multiple RDA axes. In this case the environmental mutation effect size prediction is a sum of the scores along each RDA axis, scaled by the eigenvalue of each axis and the loading of the environment along each axis. In the simulations, individual phenotypes are the sum of additive effects of mutation without dominance.

**Figure S14.**
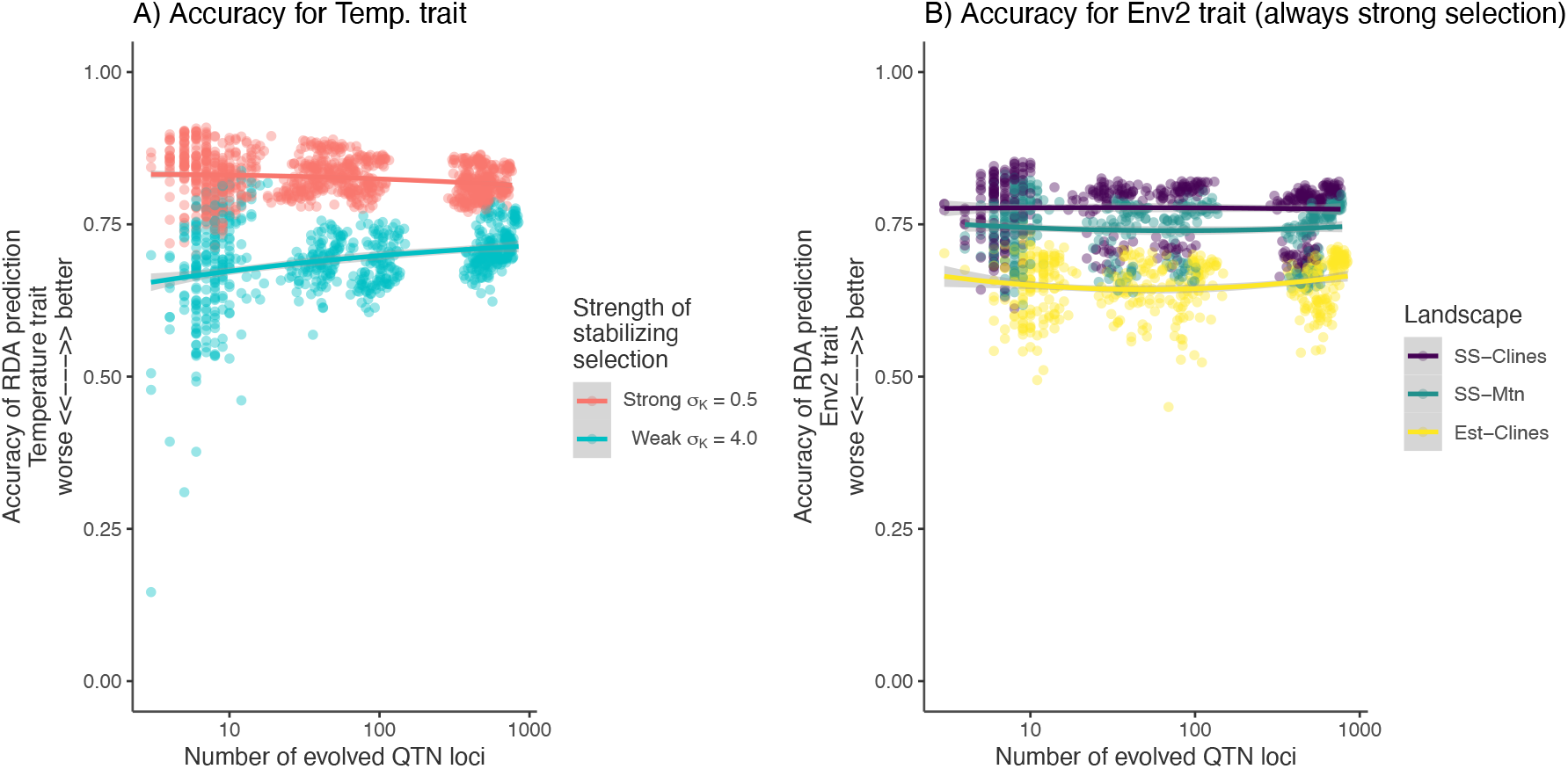
Accuracy of RDA prediction (without structure correction) as a function of the number of QTN. A) Accuracy of the RDA-predicted individual trait value for the temperature trait. The temperature trait was under selection by the same dynamics across landscapes, but was simulated under different strengths of selection. Note the higher sampling error in the oligogenic case. Weaker selection on the trait led to weaker selection on the underlying loci, and therefore less extensive linkage disequilibrium with linked neutral loci. B) Accuracy of the RDA-predicted individual trait value for the Env2 trait. The Env2 trait was always under simulated under strong selection. Abbreviations: *SS* – *Clines*: stepping stone landscape with clines in both environments; *SS* – *Mtn*: stepping stone landscape with clines in temperature environment but mountain range for Env2; *Est* – *Clines*: estuary landscape with clines in both environments.

**Figure S15.**
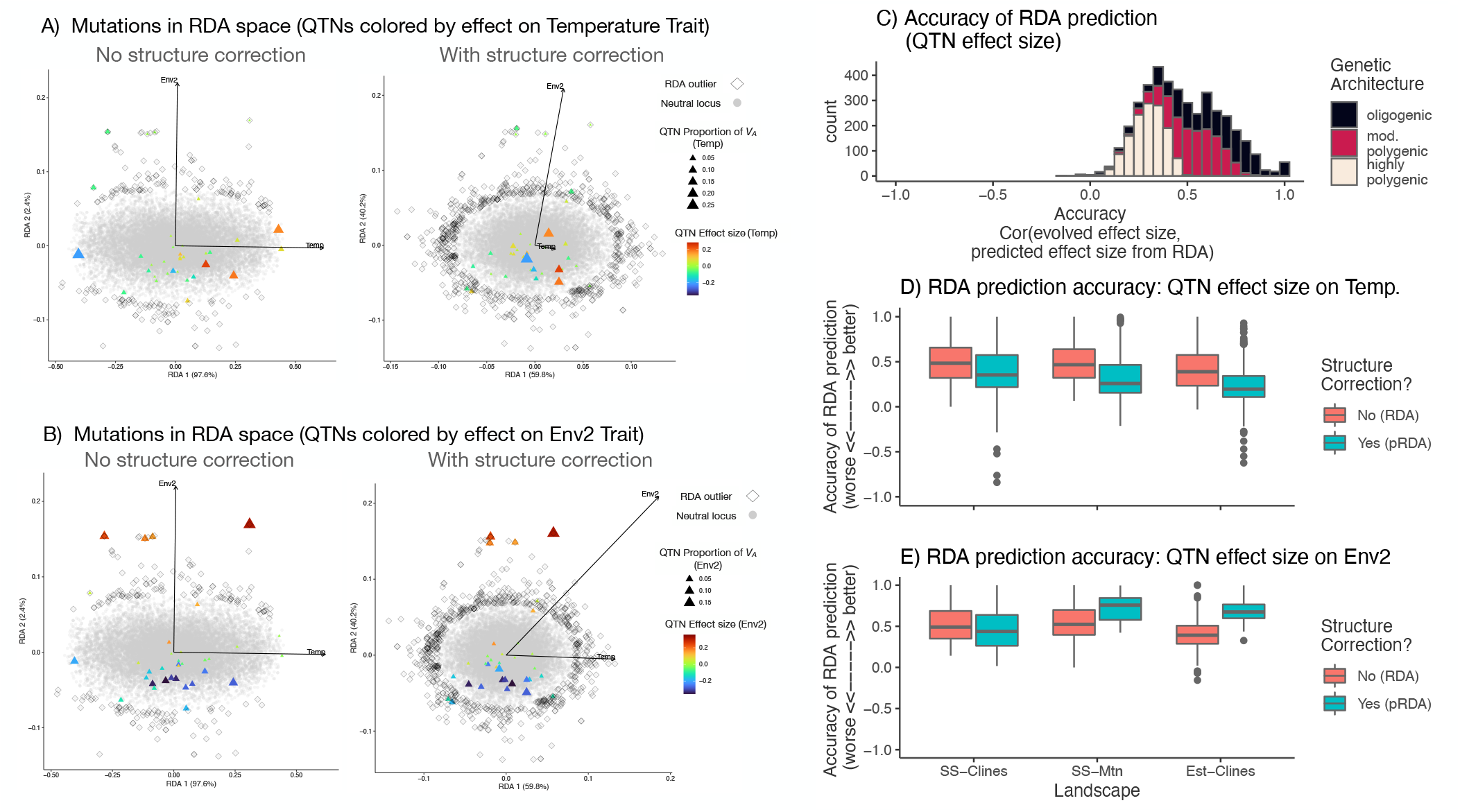
Evaluation of redundancy analysis (RDA) with and without structure correction on the mapping of QTNs. **A) and B)** Mapping of QTNs and neutral loci into RDA space for A) temperature and B) Env2. QTNs are colored according to their effect size on the trait and the point size corresponds to the proportion of additive genetic variance (*V_A_*). RDA outliers are outlined in black diamonds. Parameter levels: Estuary clines landscape, moderately polygenic, 2 traits with pleiotropy and equal selection strength, *N* central cline, *m* constant, seed 1231214. **C)** Across both traits, accuracy of the RDA-predicted QTN effect size from Eq. 1 based on its loading in RDA space without structure correction. Accuracy is measured as the correlation between the evolved QTN effect size and the RDA prediction. **D) and E)** Accuracy of the RDA-predicted QTN effect size with and without structure correction for D) mutation effects on Temperature and E) mutation effects on Env2.

**Figure S16.**
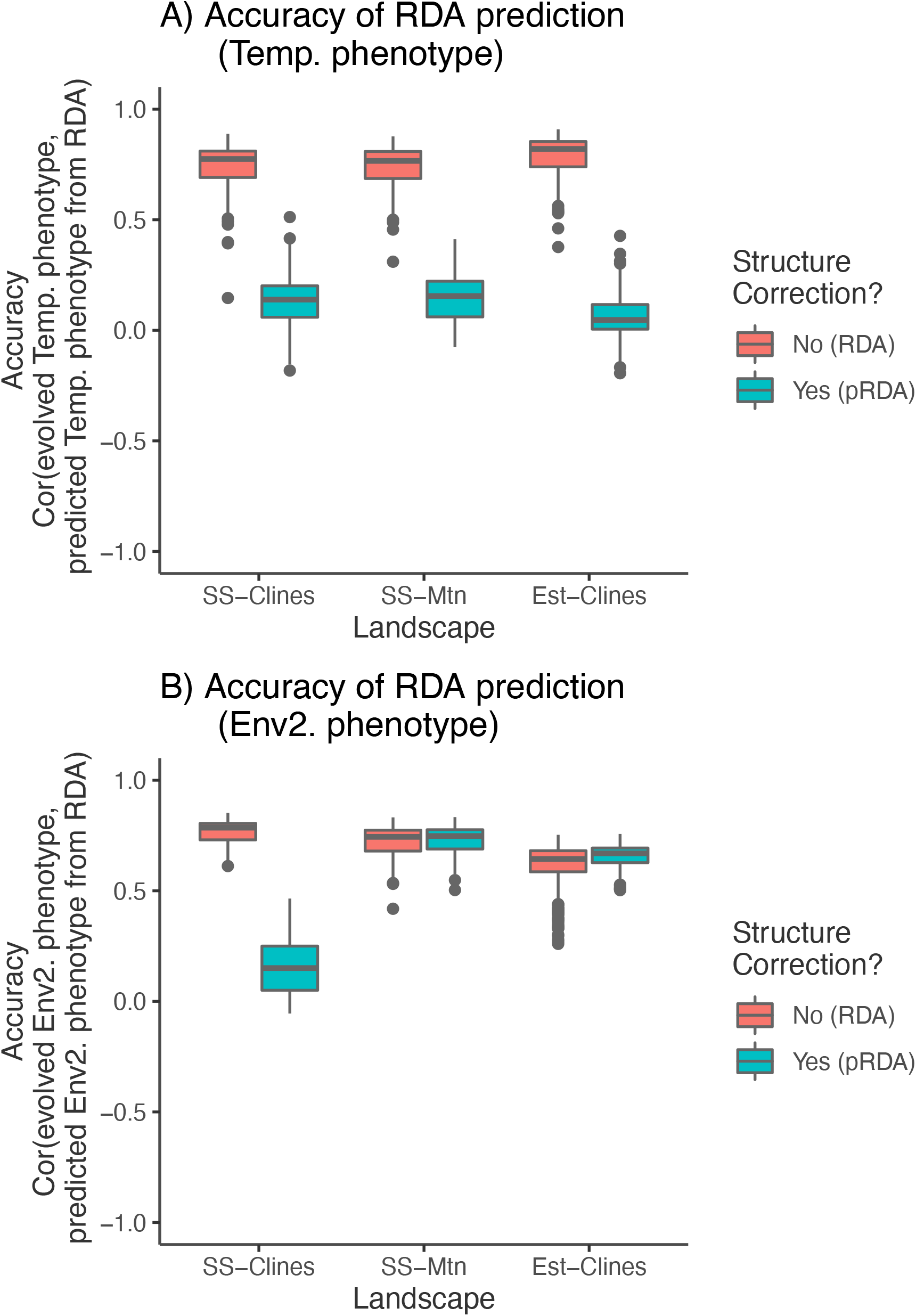
Accuracy of the RDA-predicted unscaled individual trait value with and without structure correction for A) the Temperature trait and B) Env2 trait. Accuracy is measured as the correlation between the evolved trait value and the RDA prediction for all individuals.

**Figure S17.**
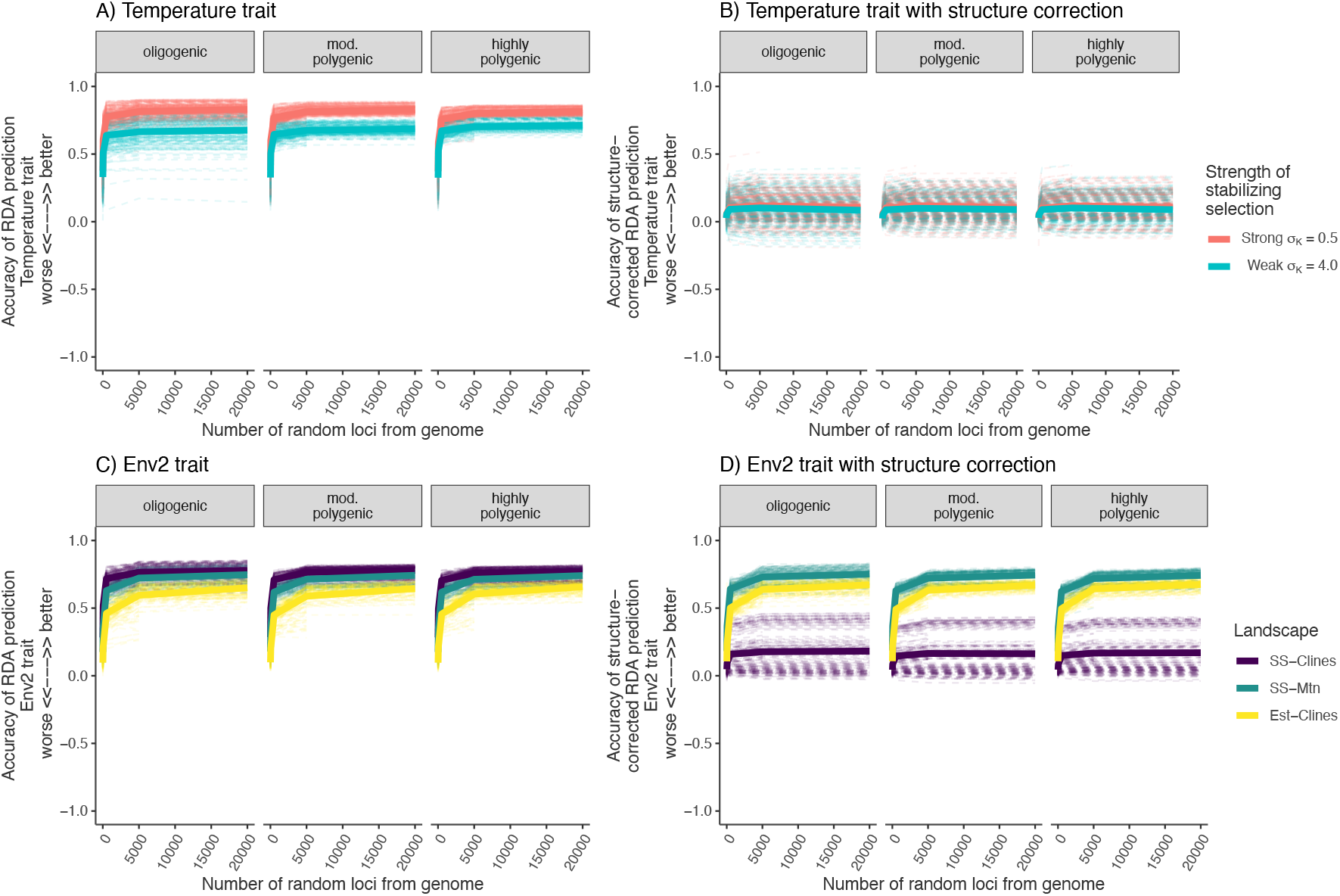
Redundancy Analysis (RDA) prediction as a function of the number of random loci sampled from the genome. Half of the genome was allowed to be affected by selection, while the other half evolved neutrally. A) Accuracy of the RDA-predicted individual trait value for the temperature trait. The temperature trait was under selection by the same dynamics across landscapes, but was simulated under different strengths of selection. Note the higher sampling error in the oligogenic case. B) Accuracy of the RDA-predicted individual trait value for the temperature trait for a partial RDA model including structure correction as the first 2 PC axes of structure. C) Accuracy of the RDA-predicted individual trait value for the Env2 trait. D) Accuracy of the RDA-predicted individual trait value for the Env2 trait for a partial RDA model including structure correction as the first 2 PC axes of structure. Abbreviations: *SS* – *Clines* Stepping Stone Clines, *SS* – *Mtn* Stepping Stone Mountain, *Est* – *Clines* Estuary Clines.

**Figure S18.**
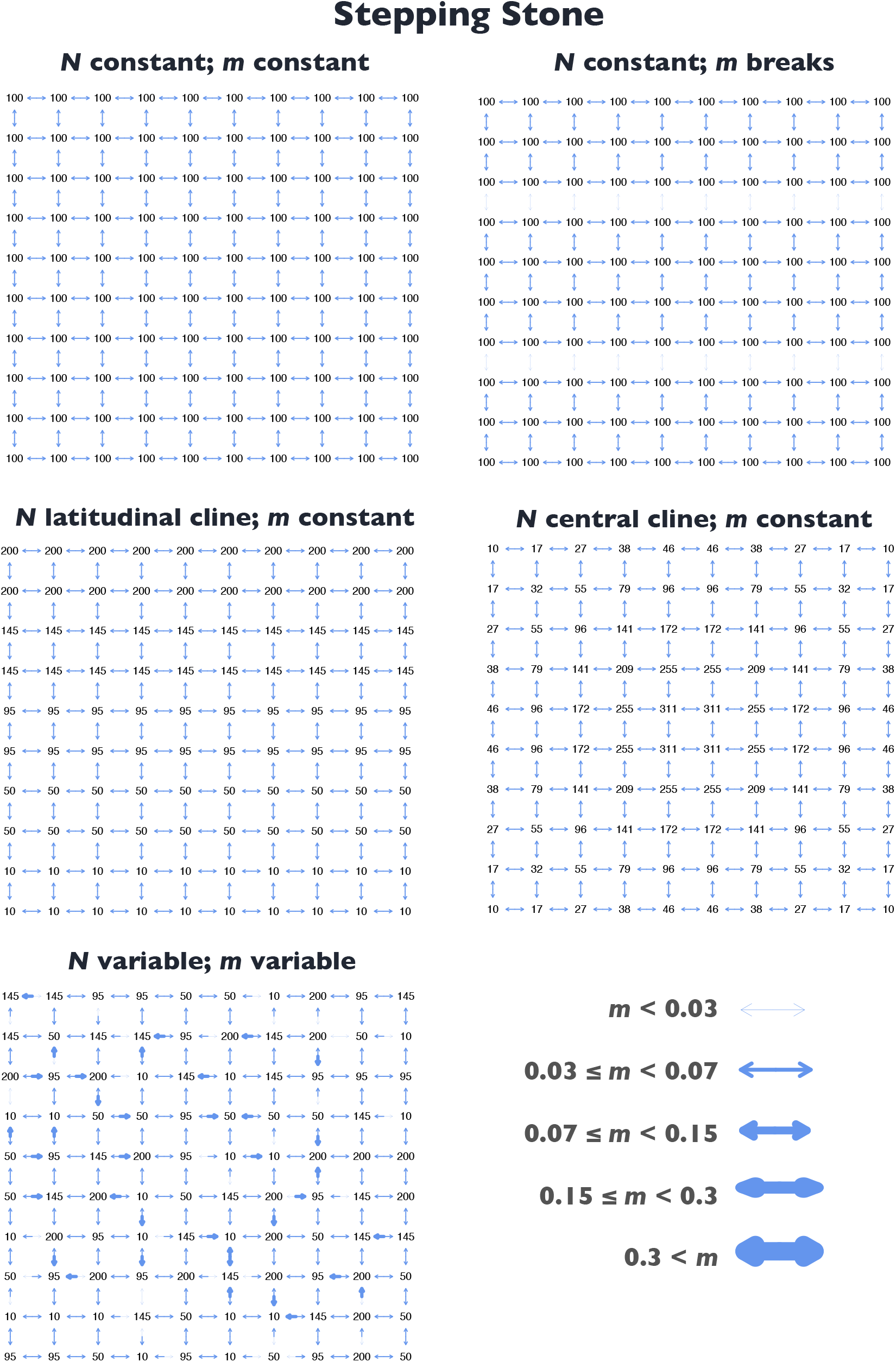
*N* and *m* for the stepping stone demographies.

**Figure S19.**
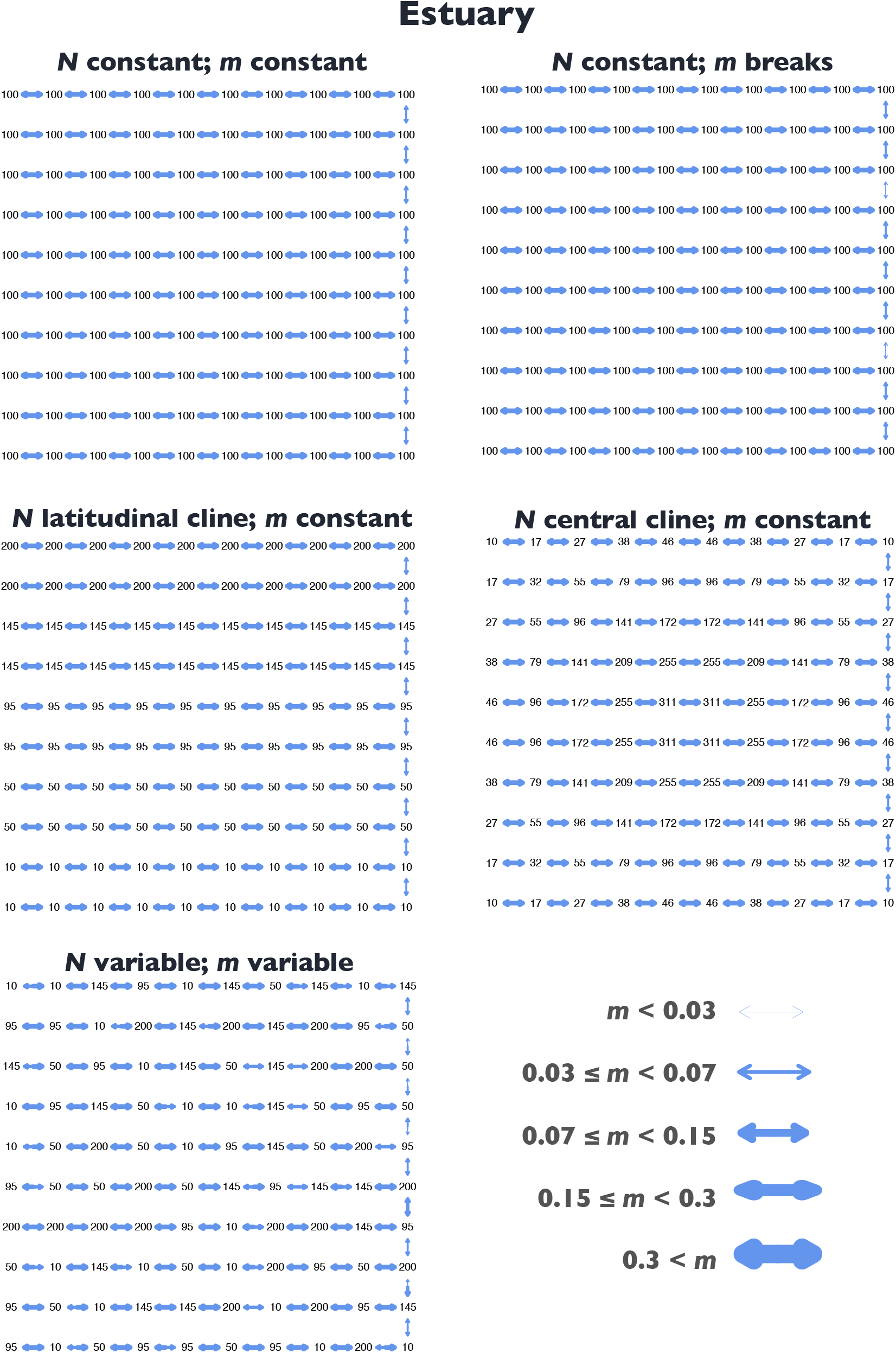
*N* and *m* for the estuary demographies.

## Literature Cited

1. Huxley, J. Clines: an auxiliary taxonomic principle. Nature 142, 219–220 (1938).

2. Endler, J.A. Geographic variation, speciation, and clines. Monogr. Popul. Biol. 10, 1–246 (1977).

3. Haldane, J. B. S. The theory of a cline. J. Genet. 48, 277–284 (1948).

4. Holgate, P. Genotype frequencies in a section of a cline. Heredity 19, 507–509 (1964).

5. May, R. M., Endler, J. A. & McMurtrie, R. E. Gene frequency clines in the presence of selection opposed by gene flow. Am. Nat. 109, 659–676 (1975).

6. Endler, J. A. Gene flow and population differentiation. Science 179, 243–250 (1973).

7. Fisher, R. A. Gene frequencies in a cline determined by selection and diffusion. Biometrics 6, 353 (1950).

8. Barton, N. H. Gene flow past a cline. Heredity 43, 333–339 (1979).

9. Dobzhansky, T. & Lewontin, R. C. Dobzhansky’s Genetics of Natural Populations I-XLIII. (Columbia University Press, 2003).

10. Walsh, B. & Lynch, M. Evolution and Selection of Quantitative Traits. (Oxford University Press, 2018).

11. Koehn, R. K., Newell, R. I. & Immermann, F. Maintenance of an aminopeptidase allele frequency cline by natural selection. Proc. Natl. Acad. Sci. U. S. A. 5385–5389 (1980).

12. Hilbish, T. J. Demographic and temporal structure of an allele frequency cline in the mussel Mytilus edulis. Mar. Biol. 86, 163–171 (1985).

13. Saavedra, C., Zapata, C., Guerra, A. & Alvarez, G. Allozyme variation in European populations of the oyster Ostrea edulis. Mar. Biol. 115, 85–95 (1993).

14. Berry, A. J. Molecular analysis of an allozyme cline: alcohol dehydrogenase in Drosophila melanogaster on the east coast of North America. Genetics 134, 869–893 (1993).

15. Holm, E. R. & Bourget, E. Selection and population genetic structure of the barnacle Semibalanus balanoides in the northwest Atlantic and Gulf of St. Lawrence. Marine Ecology Progress Series 113, 247–256(1994).

16. Rellstab, C., Gugerli, F., Eckert, A. J., Hancock, A. M. & Holderegger, R. A practical guide to environmental association analysis in landscape genomics. Mol. Ecol. 24, 4348–4370 (2015).

17. Hoban, S. et al. Finding the genomic basis of local adaptation: pitfalls, practical solutions, and future directions. Am. Nat. 188, 379–397 (2016).

18. Hedrick, P. W. Genetic polymorphism in heterogeneous environments: The Age of Genomics. Annu. Rev. Ecol. Evol. Syst. 37, 67–93 (2006).

19. Waldvogel, A.-M., Schreiber, D., Pfenninger, M. & Feldmeyer, B. Climate change genomics calls for standardized data reporting. Front. Ecol. Evol. 8,(2020).

20. Hedrick, P. W., Ginevan, M. E. & Ewing, E. P. Genetic polymorphism in heterogeneous environments. Annu. Rev. Ecol. Syst. 7, 1–32 (1976).

21. Pritchard, J. K., Pickrell, J. K. & Coop, G. The genetics of human adaptation: hard sweeps, soft sweeps, and polygenic adaptation. Current Biology 20, R208–R215 (2010).

22. Hancock, A. M., Alkorta-Aranburu, G., Witonsky, D. B. & Di Rienzo, A. Adaptations to new environments in humans: the role of subtle allele frequency shifts. Philos. Trans. R. Soc. Lond. B Biol. Sci. 365, 2459–2468 (2010).

23. Fumagalli, M. et al. Signatures of environmental genetic adaptation pinpoint pathogens as the main selective pressure through human evolution. PLoS Genet. 7, e1002355 (2011).

24. Westram, A. M., Faria, R., Johannesson, K., Butlin, R. & Barton, N. Inversions and parallel evolution. Philos. Trans. R. Soc. Lond. B Biol. Sci. 377, 20210203 (2022).

25. Barton, N. H. Clines in polygenic traits. Genet. Res. 74, 223–236 (1999).

26. Savolainen, O., Lascoux, M. & Merilä, J. Ecological genomics of local adaptation. Nat. Rev. Genet. 14, 807–820 (2013).

27. Frichot, E., Schoville, S. D., Bouchard, G. & François, O. Testing for associations between loci and environmental gradients using latent factor mixed models. Mol. Biol. Evol. 30, 1687–1699 (2013).

28. Frichot, E. & François, O. LEA: An R package for landscape and ecological association studies. Methods Ecol. Evol. 6, 925–929 (2015).

29. Günther, T. & Coop, G. Robust identification of local adaptation from allele frequencies. Genetics 195, 205–220 (2013).

30. Gautier, M. Genome-wide scan for adaptive divergence and association with population-specific covariates. Genetics 201, 1555–1579 (2015).

31. Coop, G., Witonsky, D., Di Rienzo, A. & Pritchard, J. K. Using environmental correlations to identify loci underlying local adaptation. Genetics 185, 1411–1423 (2010).

32. Caye, K., Jumentier, B., Lepeule, J. & François, O. LFMM 2: Fast and accurate inference of gene-environment associations in genome-wide studies. Mol. Biol. Evol. 36, 852–860 (2019).

33. Ahrens, C. W. et al. The search for loci under selection: trends, biases and progress. Mol. Ecol. 27, 1342–1356 (2018).

34. Lotterhos, K. E. & Whitlock, M. C. The relative power of genome scans to detect local adaptation depends on sampling design and statistical method. Mol. Ecol. 24, 1031–1046 (2015).

35. de Villemereuil, P., Frichot, E., Bazin, E., François, O. & Gaggiotti, O. E. Genome scan methods against more complex models: when and how much should we trust them? Mol. Ecol. 23, 2006–2019 (2014).

36. Forester, B. R., Lasky, J. R., Wagner, H. H. & Urban, D. L. Comparing methods for detecting multilocus adaptation with multivariate genotype-environment associations. Mol. Ecol. 27, 2215–2233 (2018).

37. Forester, B. R., Jones, M. R., Joost, S., Landguth, E. L. & Lasky, J. R. Detecting spatial genetic signatures of local adaptation in heterogeneous landscapes. Mol. Ecol. 25, 104–120 (2016).

38. Capblancq, T., Luu, K., Blum, M. G. B. & Bazin, E. Evaluation of redundancy analysis to identify signatures of local adaptation. Mol. Ecol. Resour. 18, 1223–1233 (2018).

39. Capblancq, T. & Forester, B. R. Redundancy analysis: A Swiss Army Knife for landscape genomics. Methods Ecol. Evol. 12, 2298–2309 (2021).

40. Hancock, A. M. et al. Adaptations to climate-mediated selective pressures in humans. PLoS Genet. 7, e1001375 (2011).

41. Evans, L.M. et al. Population genomics of Populus trichocarpa identifies signatures of selection and adaptive trait associations. Nat. Genet. 46, 1089–1096 (2014).

42. Yeaman, S. et al. Convergent local adaptation to climate in distantly related conifers. Science 353, 1431–1433 (2016).

43. Russell, J. et al. Exome sequencing of geographically diverse barley landraces and wild relatives gives insights into environmental adaptation. Nat. Genet. 48, 1024–1030 (2016).

44. Berg, P. R. et al. Adaptation to low salinity promotes genomic divergence in Atlantic Cod (Gadus morhua L.). Genome Biol. Evol. 7, 1644–1663 (2015).

45. Kuhn, T. S. The Structure of Scientific Revolutions: 50th Anniversary Edition. (University of Chicago Press, 2012).

46. Kuhn, T. S. in The Essential Tension: Selected Studies in Scientific Tradition and Change (ed. Suppe, F.) 293–319 (University of Chicago Press, 1974).

47. Haller, B. C. & Messer, P. W. SLiM 3: Forward genetic simulations beyond the Wright-Fisher model. Mol. Biol. Evol. 36, 632–637 (2019).

48. Haller, B. C., Galloway, J., Kelleher, J., Messer, P. W. & Ralph, P. L. Tree-sequence recording in SLiM opens new horizons for forward-time simulation of whole genomes. Mol. Ecol. Resour. 19, 552–566 (2019).

49. Ralph, P. L. & Coop, G. Convergent evolution during local adaptation to patchy landscapes. PLoS Genet. 11, el005630 (2015).

50. Láruson, A. J., Yeaman, S. & Lotterhos, K. E. The importance of genetic redundancy in evolution. Trends Ecol. Evol. (2020). doi:10.1016/j.tree.2020.04.009

51. Lasky, J. R. et al. Characterizing genomic variation of *Arabidopsis thaliana*: the roles of geography and climate. Mol. Ecol. 21, 5512–5529 (2012).

52. Rellstab, C., Dauphin, B. & Exposito-Alonso, M. Prospects and limitations of genomic offset in conservation management. Evol. Appl. 14, 1202–1212 (2021).

53. Legendre, P. & Legendre, L. Numerical ecology. Elsevier. Amsterdam, The Netherlands 853 (1998).

54. Havird, J. C. et al. Distinguishing between active plasticity due to thermal acclimation and passive plasticity due to Q 10 effects: Why methodology matters. Funct. Ecol. 34, 1015–1028 (2020).

55. Yeaman, S. Local adaptation by alleles of small effect. Am. Nat. 186 Suppl 1, S74–89 (2015).

56. Lotterhos, K. E., Yeaman, S., Degner, J., Aitken, S. & Hodgins, K. A. Modularity of genes involved in local adaptation to climate despite physical linkage. Genome Biol. 19, 157 (2018).

57. Wagner, G.P. Homologues, natural kinds and the evolution of modularity. Integr. Comp. Biol. 36, 36–43 (1996).

58. Griswold, C. K. Pleiotropic mutation, modularity and evolvability. Evol. Dev. 8, 81–93 (2006).

59. Le Nagard, H., Chao, L. & Tenaillon, O. The emergence of complexity and restricted pleiotropy in adapting networks. BMC Evol. Biol. 11, 326 (2011).

60. Lind, B. M., Menon, M., Bolte, C. E., Faske, T. M. & Eckert, A. J. The genomics of local adaptation in trees: are we out of the woods yet? Tree Genet. Genomes 14, 29 (2018).

61. Thompson, K. A. Experimental hybridization studies suggest that pleiotropic alleles commonly underlie adaptive divergence between natural populations. Am. Nat. 196, E16–E22 (2020).

62. Mahony, C.R. et al. Evaluating genomic data for management of local adaptation in a changing climate: A lodgepole pine case study. Evol. Appl. 13, 116–131 (2020).

63. Lotterhos, K. E., Láruson, A. J. & Jiang, L.-Q. Novel and disappearing climates in the global surface ocean from 1800 to 2100. Sci. Rep. 11, 15535 (2021).

64. Williams, J. W., Jackson, S. T. & Kutzbach, J. E. Projected distributions of novel and disappearing climates by 2100 AD. Proc. Natl. Acad. Sci. U.S. A. 104, 5738–5742 (2007).

65. Funk, W. C., McKay, J. K., Hohenlohe, P. A. & Allendorf, F. W. Harnessing genomics for delineating conservation units. Trends Ecol. Evol. 27, 489–496 (2012).

66. Aitken, S. N. & Whitlock, M. C. Assisted gene flow to facilitate local adaptation to climate change. Annu. Rev. Ecol. Evol. Syst. 44, 367–388 (2013).

67. Urban, M. C. et al. Improving the forecast for biodiversity under climate change. Science 353,(2016).

68. Reiskind, M. O. B., Moody, M. L., Bolnick, D. I., Hanifin, C. T. & Farrior, C. E. Nothing in evolution makes sense except in the light of biology. Bioscience (2021). doi:10.1093/biosci/biaa170

69. Waldvogel, A.-M. et al. Evolutionary genomics can improve prediction of species’ responses to climate change. Evol Lett 4, 4–18 (2020).

70. Láruson, A. J., Fitzpatrick, M. C., Keller, S. R., Haller, B. C. & Lotterhos, K. E. Seeing the forest for the trees: Assessing genetic offset predictions from gradient forest. Evol. Appl. 15, 403–416 (2022).

71. Kardos, M. & Shafer, A.B.A. The peril of gene-targeted conservation. Trends Ecol. Evol. 33, 827–839 (2018).

72. Russello, M. A., Kirk, S. L., Frazer, K. K. & Askey, P. J. Detection of outlier loci and their utility for fisheries management. Evol. Appl. 5, 39–52 (2012).

73. Flanagan, S. P., Forester, B. R., Latch, E. K., Aitken, S. N. & Hoban, S. Guidelines for planning genomic assessment and monitoring of locally adaptive variation to inform species conservation. Evol. Appl. 11, 1035–1052 (2018).

74. Collard, B.C.Y. & Mackill, D. J. Marker-assisted selection: an approach for precision plant breeding in the twenty-first century. Philos. Trans. R. Soc. Lond. B Biol. Sci. 363, 557–572 (2008).

75. Jannink, J.-L., Lorenz, A. J. & Iwata, H. Genomic selection in plant breeding: from theory to practice. Brief. Funct. Genomics 9, 166–177 (2010).

76. Heffner, E. L., Sorrells, M. E. & Jannink, J.-L. Genomic selection for crop improvement. Crop Sci. 49, 1–12 (2009).

77. Eggen, A. The development and application of genomic selection as a new breeding paradigm. Anim Fron 2, 10–15 (2012).

78. Lotterhos, K. E. The effect of neutral recombination variation on genome scans for selection. G3 9, 1851–1867 (2019).

79. Lotterhos, K. E., Fitzpatrick, M. C. & Blackmon, H. Simulation tests of methods in evolution, ecology, and systematics: pitfalls, progress, and principles. Annual Reviews of Ecology, Evolution, and Systematics in press,

80. Jones, F. C. et al. The genomic basis of adaptive evolution in threespine sticklebacks. Nature 484, 55–61 (2012).

81. Eierman, L. E. & Hare, M. P. Survival of oyster larvae in different salinities depends on source population within an estuary. J. Exp. Mar. Bio. Ecol. 449, 61–68 (2013).

82. Fisher, R. A. The Genetical Theory of Natural Selection. 302 (Oxford, Clarendon Press, 1930, 1930).

83. Wright, S. Evolution in Mendelian populations. Genetics 97–159 (1931).

84. Wright, S. The distribution of gene frequencies under irreversible mutation. Proc. Natl. Acad. Sci. U. S. A. 24, 253–259 (1938).

85. Crow, J. F. & Kimura, M. An Introduction to Population Genetics Theory. (Harper and Row, Publishers, Inc., 1970).

86. Bürger, R. The mathematical theory of selection, recombination, and mutation. (Wiley, 2000).

87. Kelleher, J., Thornton, K. R., Ashander, J. & Ralph, P. L. Efficient pedigree recording for fast population genetics simulation. PLoS Comput. Biol. 14, e1006581 (2018).

88. Danecek, P. et al. The variant call format and VCFtools. Bioinformatics 27, 2156–2158 (2011).

89. Blanquart, F., Kaltz, O., Nuismer, S. L. & Gandon, S. A practical guide to measuring local adaptation. Ecol. Lett. 16, 1195–1205 (2013).

90. Weir, B. S. & Cockerham, C. C. Estimating F-statistics for the analysis of population structure. Evolution 38, 1358–1370 (1984).

91. Whitlock, M. C. & Lotterhos, K. E. Reliable detection of loci responsible for local adaptation: inference of a null model through trimming the distribution of *F_ST_*. Am. Nat. 186, S24–S36 (2015).

92. Kendall, M. G. A new measure of rank correlation. Biometrika 30, 81–93 (1938).

93. Kendall, M. G. The treatment of ties in ranking problems. Biometrika 33, 239–251 (1945).

94. Cattell, R. B. The scree test for the number of factors. Multivariate Behav. Res. 1, 245–276 (1966).

95. Dixon, P. VEGAN, a package of R functions for community ecology. J.Veg. Sci. 14, 927–930 (2003).

96. Dabney, Storey & Warnes. qvalue: Q-value estimation for false discovery rate control. R package version

## References

Eierman, L. E., & Hare, M. P. (2013). Survival of oyster larvae in different salinities depends on source population within an estuary. Journal of Experimental Marine Biology and Ecology, 449, 61–68.

Jones, F. C., Grabherr, M. G., Chan, Y. F., Russell, P., Mauceli, E., Johnson, J., … Kingsley, D.M. (2012). The genomic basis of adaptive evolution in threespine sticklebacks. Nature, 484(1392), 55–61.

Legendre, P., & Legendre, L. (1998). Numerical ecology. Elsevier. Amsterdam, The Netherlands, 853.

